# Structural basis for RNA-mediated assembly of type V CRISPR-associated transposons

**DOI:** 10.1101/2022.06.17.496590

**Authors:** Michael Schmitz, Irma Querques, Seraina Oberli, Christelle Chanez, Martin Jinek

## Abstract

CRISPR systems have been co-opted by Tn7-like elements to direct RNA-guided transposition. Type V-K CRISPR-associated transposons rely on the concerted activities of the pseudonuclease Cas12k, the AAA+ ATPase TnsC, the Zn-finger protein TniQ, and the transposase TnsB. Here we present a cryo-electron microscopic structure of a target DNA-bound Cas12k-transposon recruitment complex comprising RNA-guided Cas12k, TniQ, TnsC and, unexpectedly, the ribosomal protein S15. Complex assembly on target DNA results in complete R-loop formation mediated by critical interactions between TniQ and the trans-activating crRNA, and is coupled with TniQ-dependent nucleation of a TnsC filament. *In vivo* transposition assays corroborate our structural findings, and biochemical and functional analyses of S15 supports its role as a *bona fide* component of the type V crRNA-guided transposition machinery. Altogether, our work uncovers key aspects of the mechanisms underpinning RNA-mediated assembly of CRISPR-associated transposons that will guide their development as programmable site-specific gene insertion tools.

## Introduction

The canonical function of prokaryotic CRISPR-Cas systems is to provide adaptive immunity against invading mobile genetic elements - including transposons, plasmids and phages. This relies on CRISPR-associated (Cas) protein effector complexes that mediate CRISPR RNA (crRNA)-guided recognition of target nucleic acids and their subsequent nucleolytic degradation (Koonin et al., 2017; Sorek et al., 2013). Several Tn7-like transposons have co-opted RNA-guided type I-F, I-B and V-K CRISPR-Cas systems to direct transposon DNA insertion into specific target sites (Faure et al., 2019; Klompe et al., 2019; Petassi et al., 2020; Peters et al., 2017; Rybarski et al., 2021; Saito et al., 2021; Strecker et al., 2019). The CRISPR modules of CRISPR-associated transposons are comprised by either a nuclease-deficient multisubunit Cascade complex in type I systems (Klompe et al., 2019) or a single catalytically inactive Cas12k pseudonuclease in type V-K systems (Strecker et al., 2019), and are encoded between the left and right transposon end sequences together with a CRISPR spacer-repeat array and several transposase genes. The DNA-targeting effectors bind to target sites specified by the crRNA guides, recruiting the transposon machinery to catalyze transposon DNA insertion at a fixed distance downstream of the specific target DNA site. As CRISPR-associated transposon (CAST) systems often contain additional defense systems as cargos (Klompe et al., 2022), these elements have been hypothesizes to mediate horizontal gene transfer of host defense systems within bacterial populations by using other transposons and plasmids as shuttle vectors. While crRNAs encoded from the CRISPR arrays guide transposon insertion preferentially into other mobile genetic elements, atypical delocalized crRNAs target integration into host chromosomal sites for transposon homing (Saito et al., 2021)

For type V-K systems, RNA-guided transposition relies on the CRISPR effector complex comprising Cas12k, a crRNA and a trans-activating RNA (tracrRNA), and three transposon proteins: the AAA+ ATPase TnsC, the transposase TnsB and the zinc-finger protein TniQ (Strecker et al., 2019). Biochemical and structural studies of a type V CAST system from *Scytonema hofmanni* (ShCAST) revealed that Cas12k-mediated DNA targeting depends on the intricate architecture of the tracrRNA that serves as a scaffold to correctly position Cas12k and the crRNA guide for the recognition of complementary targets (Park et al., 2021; Querques et al., 2021; Xiao et al., 2021). Upon binding to a 5’-GTN protospacer adjacent motif (PAM) in the target DNA, Cas12k initiates guide RNA hybridization that generates an incomplete R-loop structure, suggesting that further rearrangements occurring upon recruitment of the downstream transposon machinery are required to elicit further guide RNA-target DNA hybridization (Strecker et al., 2019). In turn, structural and biochemical studies revealed that TnsC assembles ATP-dependent helical filaments on double-stranded DNA, remodeling its duplex geometry (Park et al., 2021; Querques et al., 2021). Based on homology to the prototypical *E. coli* Tn7 transposon (Peters and Craig, 2001) and analysis of transposase-mediated integration events (Vo et al., 2021a), the DDE-type TnsB transposase has been postulated to catalyze the 3′-DNA strand breakage and transfer reactions required for a replicative transposition mechanism, resulting in transposon end nicking and ligation to a target DNA at sites located 60-66 bp downstream of the PAM (Strecker et al., 2019). TnsC is thought to recruit and activate TnsB at the target site (Peters and Craig, 2001). Concurrently, TnsB interacts with the TnsC filament to trigger its disassembly by stimulating the ATPase activity of TnsC (Park et al., 2021; Querques et al., 2021). This prevents insertion of additional transposon copies into the same site and ensures target immunity, which is a conserved feature of Tn transposons harboring coupled transposase and AAA+ ATPase components (Adzuma and Mizuuchi, 1988; Greene and Mizuuchi, 2002) (Skelding et al., 2003) (Klompe et al., 2019; Strecker et al., 2019). TnsC forms a hexameric ring when bound to dsDNA in the ADP-bound state, suggesting that this intermediate might bridge between the Cas12k-DNA targeting complex and TnsB upon ATP hydrolysis, thereby providing a molecular ruler mechanism for insertion site definition (Park et al., 2021). However, the structural details of a complete CRISPR-transposon assembly remain elusive. The function of TniQ is also presently unclear. In type V-K systems, TniQ directly interacts with one end of the TnsC filament and has been implicated in the regulation of its polymerization (Park et al., 2021; Querques et al., 2021). Conversely, in type I-F3 systems a TniQ homodimer is an integral component of the DNA targeting complex together with the Cascade (Halpin-Healy et al., 2019; Jia et al., 2020; Li et al., 2020).

CRISPR-associated transposons constitute programmable, targeted DNA integration machineries that have been repurposed as site-specific, homology-independent DNA insertion tools to engineer bacterial hosts (Vo et al., 2021b; Zhang et al., 2020) and communities (Rubin et al., 2022). However, their application in eukaryotic cells has so far been hindered, in part by limited understanding of their underlying molecular mechanisms. Here, we used cryo-electron microscopy (cryo-EM) to visualize the structure of a Cas12k-transposon recruitment complex from ShCAST (Strecker et al., 2019), revealing how guide RNA-bound Cas12k, TniQ, TnsC and, unexpectedly, the ribosomal protein S15 cooperatively assemble with each other on target DNA. The structure identifies TniQ as a key structural element that bridges Cas12k and TnsC via interactions with the tracrRNA and the PAM-distal end of a complete R-loop structure, priming the nucleation of the TnsC filament in a productive orientation. We further identify the host-encoded protein S15 as a *bona fide* component of the crRNA-guided transposition machinery that promotes complex assembly and enhances transposition activity by a structural mechanism reminiscent of its function in cellular translation. Altogether, our work elucidates how a CRISPR effector recruits its cognate transposon complex to initiate RNA-guided transposition and will inform reconstitution of this machinery for programmable site-specific DNA insertion in genome engineering applications.

## Results

### Reconstitution and structure of Cas12k-transposon recruitment complex

To obtain structural insights into the interaction between Cas12k and the transposon components in type V CRISPR-associated transposons, we sought to reconstitute guide RNA-programmed Cas12k together with TnsC and TniQ on target DNA. To this end, we first bound a single-molecule guide RNA (sgRNA), comprising sequences corresponding to the crRNA guide and a trans activating crRNA (tracrRNA), and an internally unpaired target DNA oligonucleotide duplex to Cas12k immobilized on a solid support, and subsequently incubated the resulting complex with TnsC and TniQ in the presence ATP to trigger TnsC filament assembly. After extensive washing and elution, the sample was vitrified and imaged using cryo-EM for single-particle analysis. By stringent selection of Cas12k particles displaying adjacent filament-like densities, and subsequent 2-D/3-D classification, we were able to obtain a reconstruction of the Cas12k-transposon recruitment complex at a resolution of 3.3 Å (**Figure 1, Figure S1, Table S1**).

**Figure 1.**
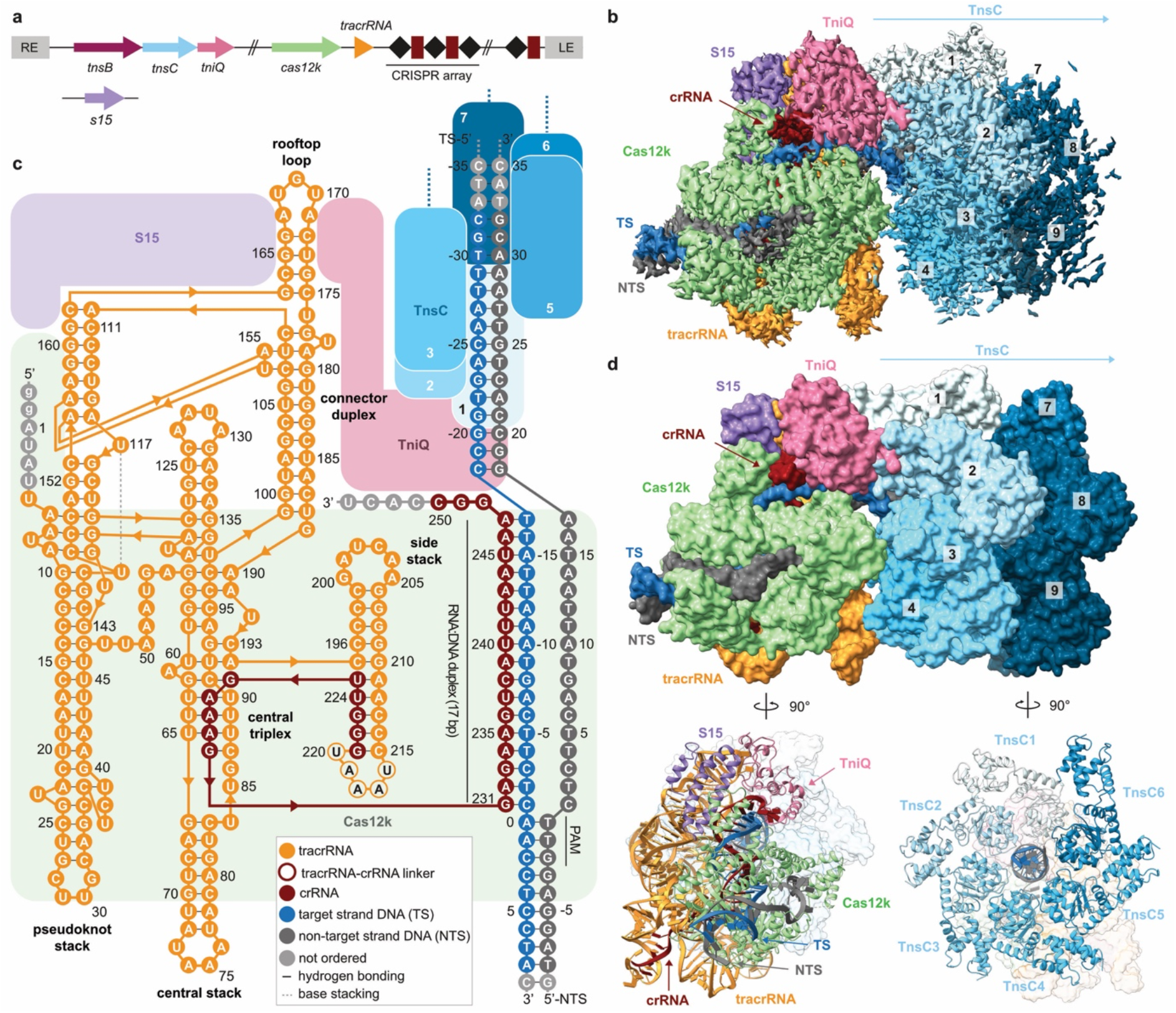
Cryo-EM structure of the Cas12k-TnsC transposon recruitment complex. (**A**) Schematic of the type V-K CRISPR-associated transposon system from *Scytonema hofmanni* (Strecker et al., 2019) and the S15 gene. LE, RE: left and right transposon ends. (**B**) Cryo-EM density map of the Cas12k-transposon recruitment complex resonctruction. Density for nine TnsC protomers (TnsC1-TnsC9) is shown. (**C**) Schematic of the sgRNA structure and R-loop architecture indicating interactions between nucleic acids and protein components of the complex. (**D**) Structural model of the Cas12k-transposon recruitment complex (surface and cartoon representations). TS, target DNA strand; NTS, non-target DNA strand.

The resulting atomic model of the complex comprises Cas12k and the sgRNA guide bound to the target site in the DNA, together with a single TniQ molecule and an emerging right-handed TnsC filament assembled on the PAM-distal region of the target DNA. Cas12k binds the DNA in a guide RNA- and PAM-dependent manner, as observed previously in the structure of the Cas12k-sgRNA-target DNA complex (Querques et al., 2021; Xiao et al., 2021). TniQ makes direct contacts with the tracrRNA part of the sgRNA and two TnsC protomers, thereby bridging Cas12k and the TnsC filament without directly interacting with Cas12k. The TnsC filament is assembled with the TnsC C-terminal domains pointing away from Cas12k. The reconstruction contains additional proteinaceous density, which we were able to assign to a single copy of the *Escherichia coli* ribosomal S15 protein that was serendipitously co-purified with TniQ, as verified by mass-spectrometric analysis of the TniQ sample (**Table S2**). Notably, S15 makes extensive contacts with both Cas12k and the tracrRNA part of the sgRNA (**Figure 1C**,**D**).

Within the same cryo-EM sample, we were additionally able to identify a separate population of particles comprising Cas12k bound to DNA together with TnsC oligomers assembled in the reverse orientation, i.e. with TnsC C-terminal domains pointing towards Cas12k, obtaining a reconstruction at a resolution of 4.1 Å (**Figure S2**). Notably, the R-loop in this assembly remained in its incomplete form and both TniQ and S15 are absent (**Figure S3**). Furthermore, there are no direct intermolecular contacts between Cas12k-sgRNA and the proximal end of the TnsC filament, suggesting that this molecular assembly represents a non-productive complex in which TnsC spontaneously oligomerized on the target DNA in a Cas12k-independent manner.

### R-loop completion occurs upon TniQ and TnsC binding

Previous structures of the Cas12k-guide RNA-target DNA complexes revealed incomplete hybridization of the crRNA sequence and the target strand (TS) of the DNA, resulting in a nine-base pair (bp) duplex (Querques et al., 2021; Xiao et al., 2021). In the present structure of the Cas12k-transposon recruitment complex, the crRNA and the target DNA form a complete R-loop structure comprising 17 base pairs, beyond which the TS and the non-target strand (NTS) rehybridize. crRNA-TS DNA hybridization beyond the 17^th^ base pair is prevented by TniQ binding to the complete R-loop, leaving seven unpaired nucleotides at the 3’ end of the crRNA spacer sequence. This observation is consistent with previous studies showing that 3’-terminally truncated crRNAs comprising 17-nucleotide spacer segments supported type V CRISPR-associated transposon activity *in vivo* (Saito et al., 2021). Overall, the DNA adopts a bent conformation, with the PAM-distal DNA exiting Cas12k at a 122° angle relative to the PAM-proximal DNA duplex (**Figure 2A**). The backbone of the displaced NTS can be completely traced as it wraps around Cas12k, passing through a gap between the REC lobe and the RuvC domain (**Figure 2A**). Completion of the R-loop occurs by TniQ interacting with the PAM-distal end of the sgRNA-TS heteroduplex, and is enabled by conformational rearrangements within Cas12k (**Figure 2A**,**B**). The Cas12k bridge helix, which precludes full R-loop formation in the structure of the Cas12k-sgRNA-target DNA complex (Querques et al., 2021; Xiao et al., 2021), is repositioned to expose the binding cleft for the PAM-distal part of the sgRNA-TS DNA heteroduplex (**Figure S4A**,**B**). This is accompanied by structural ordering of the REC lobe motifs, whereby the REC1 domain (residues 12-239^Cas12k^) interacts with the unpaired NTS while the REC2 domain (residues 240-278^Cas12k^) contacts the extended crRNA-TS heteroduplex. Further conformational rearrangements occur in the RuvC domain, where an alpha-helical hairpin (residues 548-590^Cas12k^) is repositioned to contact the NTS and the tracrRNA scaffold (**Figure S4B**,**C**).

**Figure 2.**
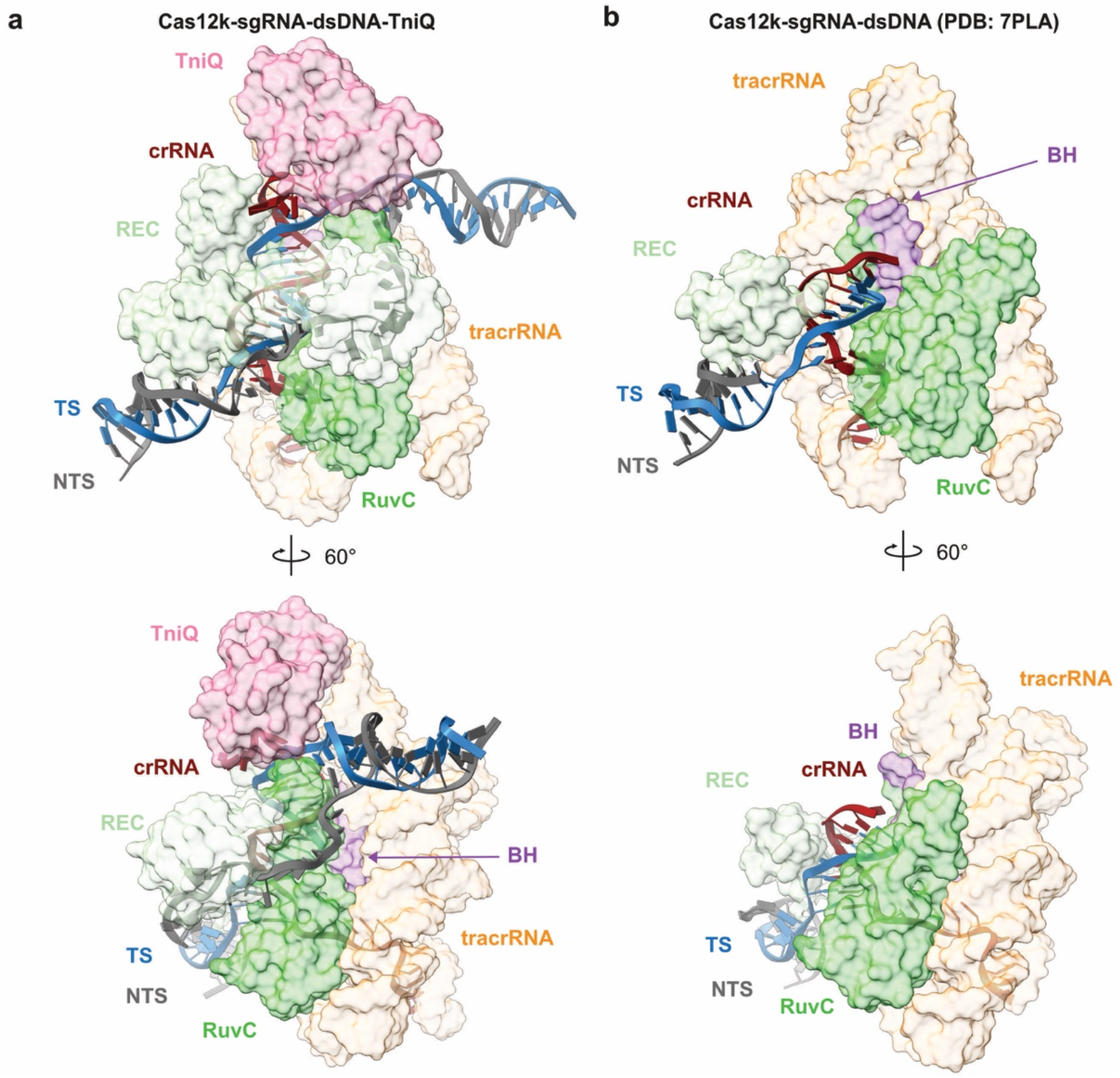
R-loop completion upon complex assembly. (**A**) Detailed views of the R-loop structure in the Cas12k-transposon recruitment complex, comprising the crRNA portion of the single guide RNA (red cartoon backbone), the target DNA strand (TS, blue cartoon backbone) and the non-target DNA strand (NTS, dark grey cartoon backbone). Only the REC (recognition lobe) and RuvC domains and the bridging helix (BH) of Cas12k are shown in surface representation. (**B**) Detailed views of the R-loop structure in the Cas12k-sgRNA-target DNA complex (PDB ID:7PLA, (Querques et al., 2021).

### TniQ recognizes tracrRNA and R-loop

In the Cas12k-transposon recruitment complex, TniQ is confined by the tracrRNA rooftop loop and the PAM-distal end of the crRNA-TS DNA heteroduplex on one side and the TnsC filament on the other (**Figure 3A**). The rooftop loop (nucleotides 167-171^tracrRNA^), which is structurally disordered in the Cas12k-sgRNA-target DNA complex (Querques et al., 2021; Xiao et al., 2021), now assumes a well-defined pentaloop conformation whose shape is read out by hydrogen bonding contacts with side chains of Gln93^TniQ^, Arg98^TniQ^, Lys128^TniQ^, Lys132^TniQ^, Gln137^TniQ^, in addition to a π-π stacking interaction between rA169^tracrRNA^ and Trp120^TniQ^ (**Figure 3B**). To validate the observed interactions, we tested the effect of TniQ and tracrRNA mutations on the transposition activity of ShCAST *in vivo* using quantitative droplet-digital PCR analysis. Mutations of a subset of interacting residues substantially reduced transposition activity. In turn, substitution of the rooftop pentaloop with a GAAA tetraloop or adenine substitution of U168^tracrRNA^ led to complete loss of transposition, while individual substitutions of other pentaloop nucleotides substantially reduced transposition activity (**Figure 3C**). TniQ further interacts with the PAM-distal end of the R-loop. Here, the terminal base pair of the crRNA-TS heteroduplex is contacted by Asn59^TniQ^ at the minor groove edge and capped by a π-π stacking interaction with His57^TniQ^, which is in turn hydrogen bonded to His94^TniQ^ (**Figure 3D**). Mutations of TniQ residues directly interacting with the crRNA-TS DNA duplex substantially reduced transposition activity *in vivo* (**Figure 3E**). Together with our structural observations, these results confirm the critical role of the tracrRNA rooftop loop as a TniQ interaction site, and its significance for TnsC recruitment to support transposition activity of type V CRISPR-associated transposon. Furthermore, the observed interactions of TniQ with the PAM-distal end of the guide RNA-TS DNA heteroduplex suggest that R-loop completion is facilitated by TniQ recruitment.

**Figure 3.**
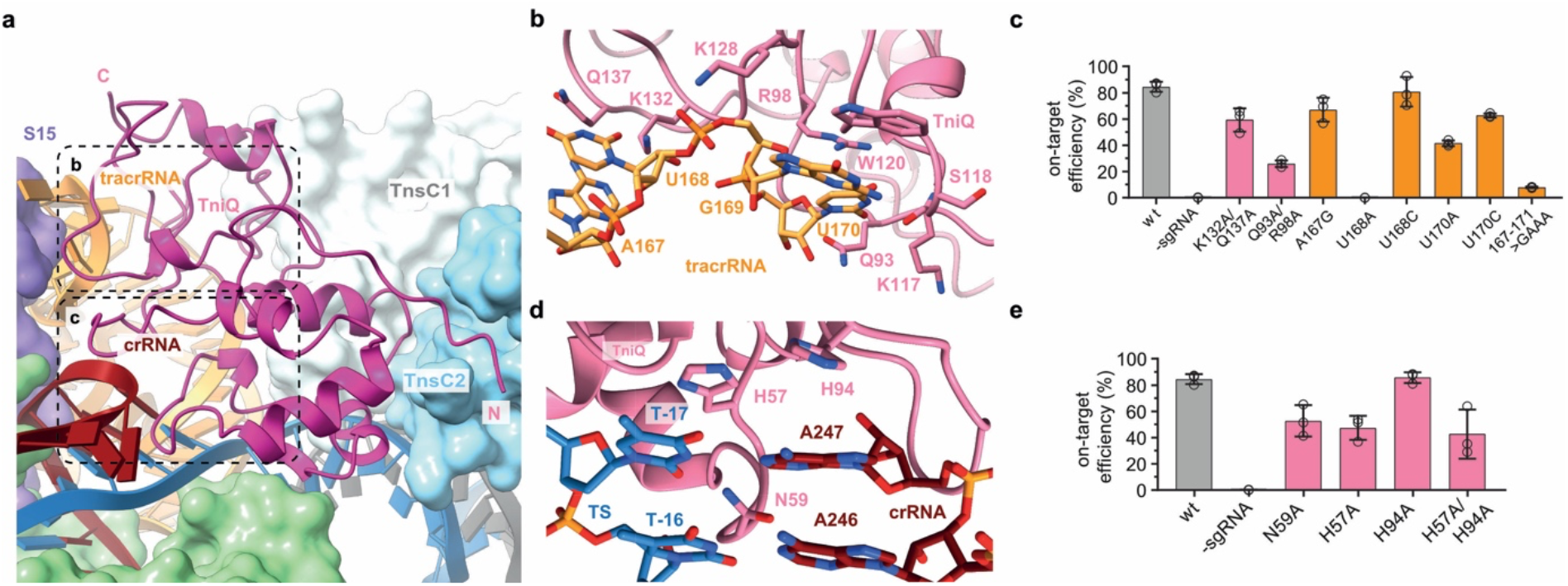
TniQ recognizes tracrRNA and completed R-loop. (**A**) Overview of the TniQ in the Cas12k-transposon recruitment complex, depicting interfaces with tracrRNA scaffold (yellow) and the RNA:DNA heteroduplex formed by the crRNA (red cartoon backbone) and the target DNA strand (TS, blue cartoon backbone). NTS (non-target DNA strand, dark grey cartoon backbone). The N- and C-termini of TniQ are indicated. (**B**) Close-up views of key tracrRNA-interacting residues of TniQ. (**C**) Site-specific transposition activity in *E. coli* of structure-based tracrRNA mutants and mutants at the tracrRNA-interacting interface, as determined by droplet digital PCR (ddPCR) analysis. Data are presented as mean ± s.d. (*n*=3 biologically independent replicates). (**D**) Detailed views of R-loop recognition by TniQ. (**E**) Site-specific transposition activity in *E. coli* of ShCAST systems structure-based tracrRNA mutants and mutants in the R-loop recognition interface of TniQ. Data are presented as mean ± s.d. (*n*=3 biologically independent replicates.

### TniQ nucleates TnsC filament formation

Positioned by interactions with the tracrRNA rooftop loop and the R-loop, the single TniQ molecule in the Cas12k-transposon recruitment complex straddles two TnsC protomers at the Cas12k-proximal end of the TnsC filament (**Figure 4A**). The C-terminal zinc finger domain (ZnF2) of TniQ contacts the terminal TnsC protomer (TnsC1), mostly via electrostatic interactions (**Figure 4B**). In turn, the N-terminal HTH domain of TniQ interacts extensively with the next TnsC protomer (TnsC2) in the filament. Notably, the N-terminal tail of TniQ inserts into a cleft in the α/β AAA+ domain of TnsC2, with the aromatic side chain of Trp10^TniQ^ sandwiched by hydrophobic interactions with Tyr115^TnsC2^ and Pro86^TnsC2^ (**Figure 4C**). Glutamate substitution of TnsC1-interacting residue Arg155^TniQ^ reduced *in vivo* transposition by ∼50%, suggesting that the interaction of TniQ with TnsC1 contributes to transposon recruitment. In contrast, N-terminal truncation of TniQ to remove residues 1-12 resulted in complete loss of transposition, while alanine substitution of Trp10^TniQ^ resulted in >90% reduction (**Figure 4D**). Together, these results validate the critical role of the TniQ N-terminal tail for the interaction with the TnsC filament and suggest its involvement in filament nucleation.

**Figure 4.**
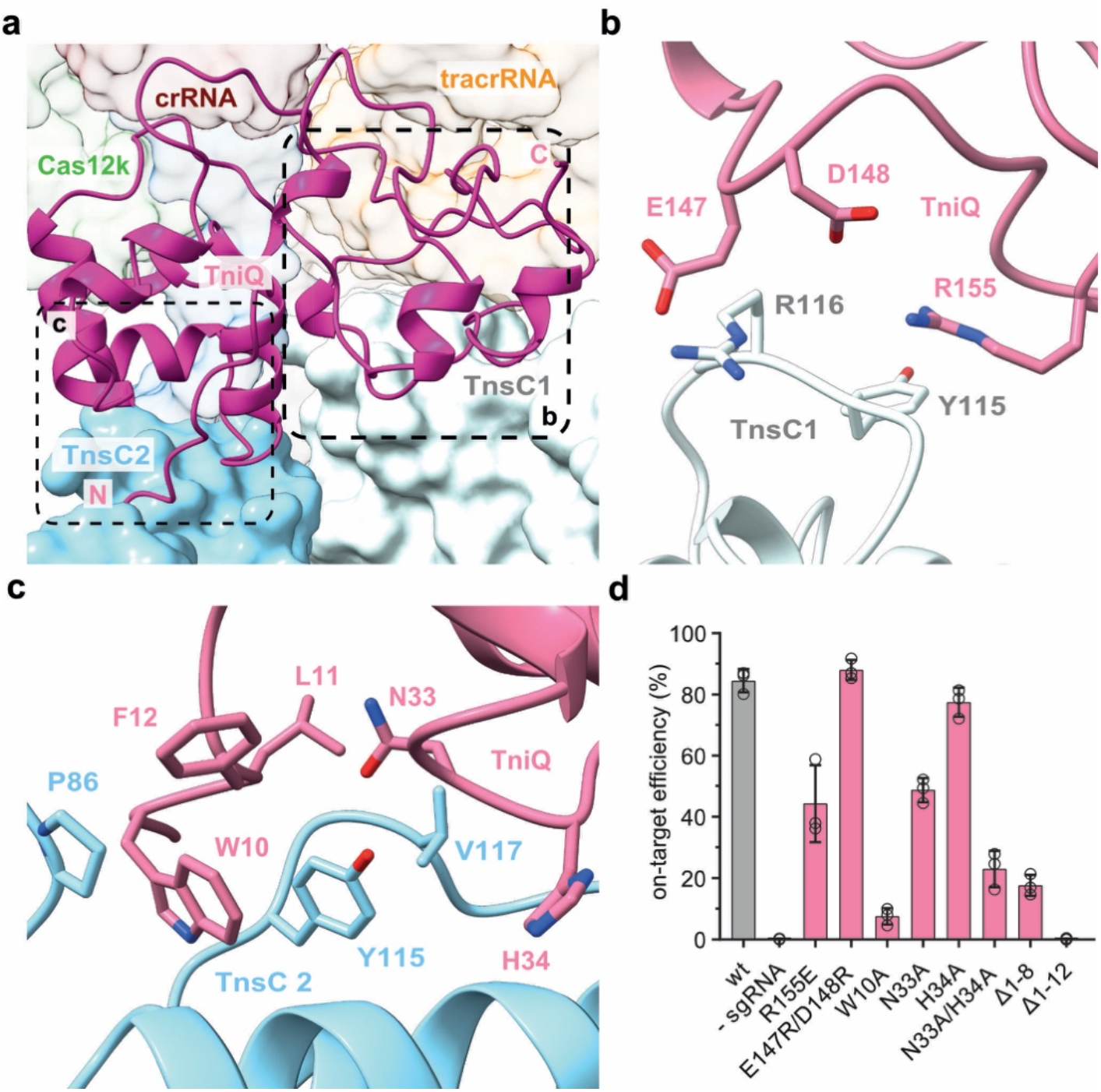
TniQ primes TnsC oligomerization. (**A**) Overview of the interactions between TniQ (cartoon representation) and two Cas12k-proximal protomers of the TnsC filament (TnsC1 and TnsC2, surface representation). The N- and C-termini of TniQ are indicated. (**B**) Close-up views of key interactions between TniQ and TnsC1. (**C**) Close up view of the interactions of TniQ with TnsC. (**D**) Site-specific transposition activity in *E. coli* of ShCAST systems containing mutations in the TnsC binding interface of TnsC, as determined by droplet digital PCR (ddPCR) analysis. Data are presented as mean ± s.d. (*n*=3 biologically independent replicates).

Previous studies have shown that in the absence of Cas12k and guide RNA, TniQ caps TnsC filaments assembled on dsDNA (Park et al., 2021), thereby restricting TnsC polymerization *in vitro* (Park et al., 2021; Querques et al., 2021). We extended these findings by docking a previously determined crystal structure of TniQ (Querques et al., 2021) into a 3.5 Å-resolution cryo-EM reconstruction of a TniQ-capped TnsC filament obtained by single-particle analysis (**Figure S5, TableS1**). Altogether, three copies of TniQ assemble at the end of the TnsC filament, each bridging a TnsC dimer within the terminal hexameric helical turn of the filament. Overall, the interactions between TnsC dimers and TniQ are highly similar with that observed in the Cas12k-transposon recruitment complex. However, the binding of Cas12k-gRNA to a TnsC filament fully capped by TniQ would not be compatible due to steric clashes with two of the three TniQ copies (**Figure S6**).

### TnsC assembles at distal end of TniQ-bound R-loop

At the PAM-distal end of the R-loop within the Cas12k-transposon recruitment complex, the TS bends away from the crRNA-TS heteroduplex and exits through a narrow channel formed by the Cas12k RuvC domain and TniQ to immediately rehybridize with the NTS (**Figure 5A**). The first base pair of the reformed TS-NTS duplex (position 18) stacks against the aromatic side chains of Tyr570^Cas12k^ and Phe567^Cas12k^, while TS nucleotides at positions 19 and 20 make backbone interactions with TniQ by hydrogen bonding with Ser36^TniQ^ and Ser38^TniQ^ (**Figure 5B**). This positions the TS-NTS duplex for entry into the TnsC helical filament. The terminal TnsC protomer (TnsC1) interacts with NTS nucleotides at positions 22 and 23 via Thr121^TnsC1^, Lys103^TnsC1^ and Lys150^TnsC1^ (**Figure 5C**). The next TnsC protomer (TnsC2) interacts with the minor groove of the duplex, contacting both TS and NTS, while TnsC3 and subsequent protomers interact mostly with the NTS. Overall, the interactions of the TnsC filament with the DNA involve the same residues (Thr121^TnsC^, Lys103^TnsC^ and Lys150^TnsC^) as previously observed in the structures of dsDNA-bound TnsC filaments and validated by mutational analysis *in vivo* (Park et al., 2021; Querques et al., 2021). Similarly, the PAM-distal DNA duplex is distorted from the canonical B-form geometry to match the helical symmetry of the TnsC filament. Crucially, the TnsC filament in the Cas12k-TnsC recruitment complex tracks the DNA strand with the opposed polarity, i.e. the NTS, as compared with the TnsC-only filament (**Figure 5D, Figure S7**).

**Figure 5.**
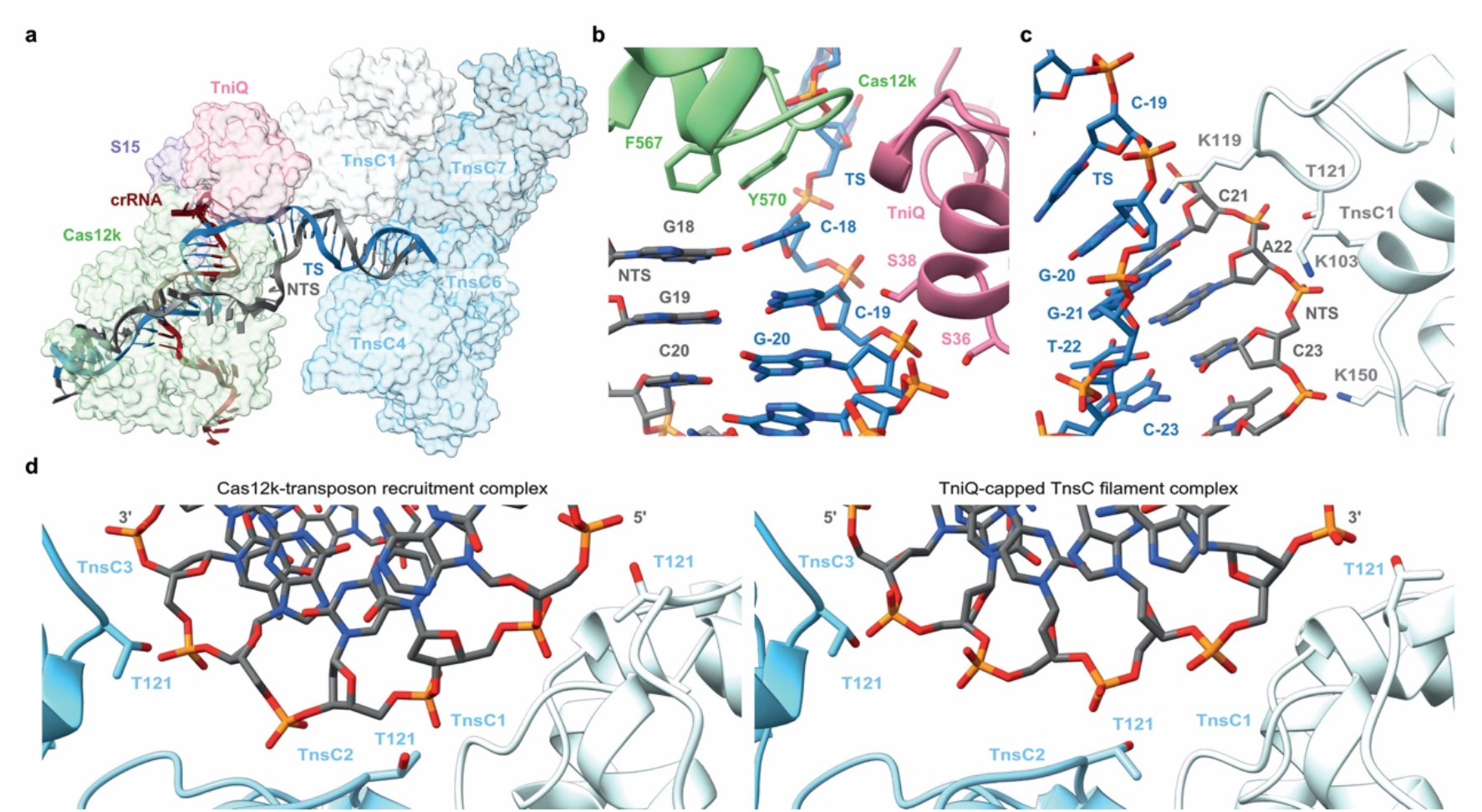
TnsC assembly on PAM distal end of R-loop DNA. (**A**) Overview of guide RNA-target DNA R-loop structure within the Cas12-transposon recruitment complex. Target strand (TS, blue cartoon backbone) and non-target DNA strand (NTS, dark grey cartoon backbone) are shown in cartoon format. Only the crRNA portion of the single guide RNA (red cartoon backbone) is shown. Proteins are shown in surface representation. Residues 132-254 of Cas12k and the TnsC2 and TnsC3 protomers are omitted from view for clarity. (**B**) Zoomed-in view of target DNA binding residues of TniQ. (**C**) Zoomed-in view of the DNA binding residues in TnsC1 protomer. (**D**) Comparison of DNA binding modes of consecutive TnsC protomers (TnsC1-TnsC3) in the Cas12k-transposon recruitment complex (left) and in the TniQ-capped TnsC filament (right).

### Ribosomal S15 protein supports Cas12k-transposon complex assembly

The *E. coli* ribosomal protein S15 (EcSh15) captured in the Cas12k-transposon recruitment complex is wedged between the Cas12k REC2 domain and the tracrRNA connector duplex and contacts the ribose-phosphate backbone of the crRNA in the PAM-distal part of the crRNA-TS DNA heteroduplex (**Fig. 6A, Figure S8A-C**). As within the small ribosomal subunit, EcS15 adopts a four-helix bundle fold and interacts with the tracrRNA and the heteroduplex in a manner that mimics its interactions with 18S rRNA (**Figure S8E**). The combined fold of EcS15 and the Cas12k REC2 domain is highly similar to the helical fold of the REC2 (Helical II) domain in the distantly related Cas12e (CasX) nuclease (Liu et al., 2019) (**Figure S8D**). EcS15 makes extensive shape-complementary interactions with the tracrRNA rooftop loop via electrostatic interactions mediated by Arg72^EcS15^, Lys73^EcS15^ and Arg77^EcS15^, and π-π stacking between Tyr69^EcS15^ and rA171^tracrRNA^, suggesting that it stabilizes the tracrRNA rooftop loop in a conformation that supports TniQ recruitment. Notably, *E. coli* and *S. hofmanni* S15 (ShS15) proteons share 58% sequence identity and the residues involved in contacting the tracrRNA and Cas12k are nearly invariant between the orthologs (**Fig S8C**). This suggests that ShS15 might act as a *bona fide* component of the transposon recruitment complex to promote the interactions of Cas12k-bound tracrRNA with TniQ, thereby contributing to TnsC recruitment. To test this hypothesis, we used pull-down experiments to observe the co-precipitation of TniQ and TnsC with immobilized StrepII-tagged Cas12k-sgRNA-target DNA complex in the presence of purified EcS15 or ShS15 proteins (**Figure 6B**). For these experiments, TniQ was re-purified according to a stringent protocol that eliminated EcS15 contamination (**Table S2**). Both EcS15 and ShS15 were efficiently co-precipitated by the Cas12k-sgRNA-DNA complex. TniQ co-precipitation was markedly enhanced in the presence of EcS15 or ShE15 and TnsC, suggesting that S15 proteins facilitate the cooperative assembly of TniQ with TnsC and Cas12k-sgRNA on target DNA. To further test the effect of EcS15 and ShS15 proteins on Cas12k-dependent transposition, we performed *in vitro* transposition assays using donor and target plasmids, sgRNA and purified recombinant proteins, monitoring transposition efficiency by droplet-digital PCR analysis. In the absence of S15 proteins, only very low levels of sgRNA-dependent transposition could be detected. Transposition efficiency was substantially enhanced by the addition of EcS15 or ShS15 (**Figure 6C**). These results indicate that S15 promotes transposition by facilitating the assembly of the Cas12k-transposon recruitment complex, suggesting that it may function as an integral part of the type V CRISPR-associated transposon machinery.

**Figure 6.**
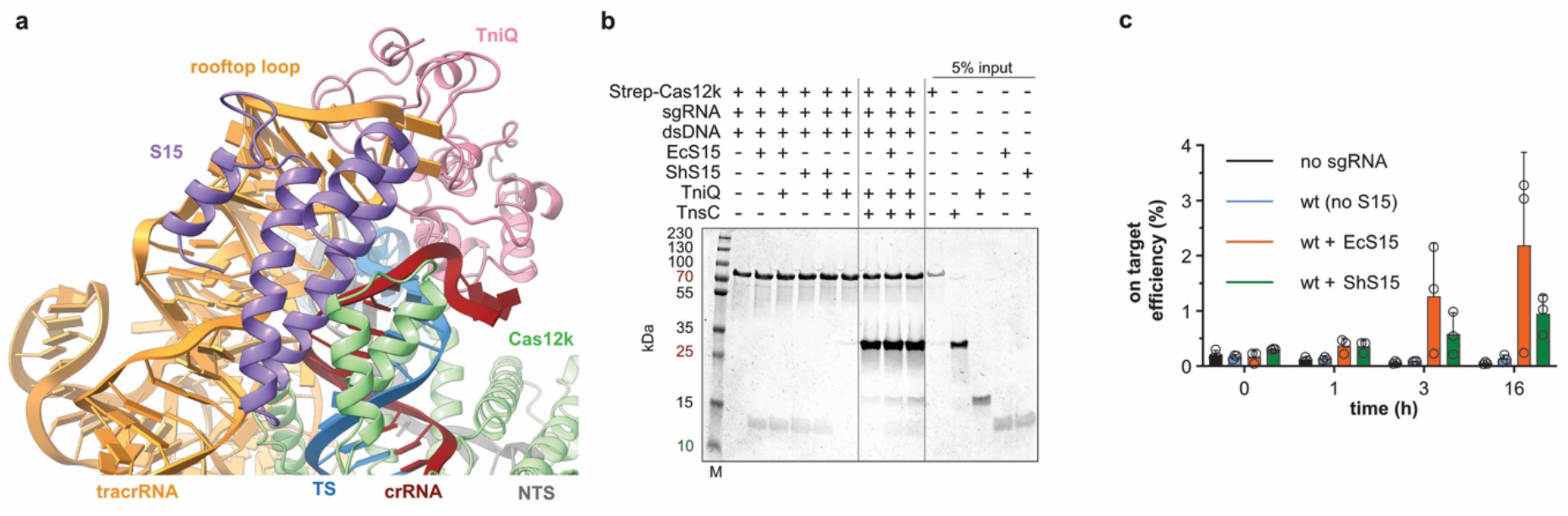
S15 promotes Cas12k-transposon recruitment complex assembly. (**A**) Zoomed-in view of S15 binding in the Cas12k-recruitment complex. (**B**) Co-precipitation of TnsC and TniQ in presence or absence of ShS15 (*Scytonema hofmanni* S15) or EcS15 (*Escherichia coli* S15) by immobilized StrepII-fused Cas12k-sgRNA-target DNA complex. (**C**) *In vitro* transposition activity of purified ShCAST components in the absence or presence of *E. coli* and *S. hofmanni* ribosomal S15 proteins, as determined by droplet digital PCR (ddPCR) analysis. Data are presented as mean ± s.d. (*n*=3 independent replicates)

## Discussion

The molecular function of CRISPR-associated transposons relies on the concerted activities of the RNA-guided CRISPR effector and transposase modules. In type V CRISPR-associated transposons, this is thought to involve interactions of the AAA+ ATPase transposon regulator TnsC with the RNA-guided pseudonuclease Cas12k at the target site but the mechanistic details have remained elusive thus far. Our structural analysis of the Cas12k-transposon recruitment complex shows that the tracrRNA component of the Cas12k guide RNA and TniQ play key roles in the process by bridging Cas12k and the ATP-dependent TnsC filament assembled on the target DNA. Our findings provide evidence that the formation of a complete R-loop structure within Cas12k occurs upon TniQ and TnsC recruitment, and thus serves as a structural checkpoint for transposon recruitment complex assembly, which is likely to have mechanistic parallels in the type I CRISPR-associated transposon systems, as hinted by previous structural analysis of the Type I-F Cascade-TniQ complex (Halpin-Healy et al., 2019; Jia et al., 2020; Li et al., 2020).

TniQ was previously shown to cap TnsC filaments and restrict their polymerization on free linear dsDNA *in vitro* (Park et al., 2021; Querques et al., 2021). Based on these observations, we proposed a mechanistic model in which we placed TniQ at the Cas12k-distal end of the TnsC filament, restricting its polymerization to the vicinity of Cas12k (Querques et al., 2021). An alternative model posited that TnsC polymerization initiates randomly on DNA and is selectively stabilized by interactions with target site-bound Cas12k and TniQ (Park et al., 2021). Our Cas12k-transposon recruitment complex structure reveals that a single TniQ copy bridges two TnsC protomers at the Cas12k-proximal end of the TnsC filament. Based on these findings and their functional validation, we thus propose a revised model in which TniQ neither caps the distal ends of TnsC filaments nor stabilizes filaments randomly nucleated on DNA (**Figure 7**). Instead, the cooperative assembly of Cas12k, guide RNA and TniQ directly nucleates TnsC filament formation starting at the PAM-distal end of the Cas12k R-loop. Although our structural and functional data do not rule out a mechanism based on randomly-initiated TnsC polymerization followed by specific capture and stabilization by DNA-bound Cas12k (Park et al., 2021), a parsimonious interpretation points to site-specific TnsC filament nucleation at the target site defined by the Cas12k guide RNA. This is also supported by the observation that the polarity of the tracking DNA strand on which the TnsC filament assembles in the context of the Cas12k-transposon recruitment complex (the NTS strand) appears to be opposite to that observed in standalone TnsC filaments assembled on free dsDNA (Park et al., 2021; Querques et al., 2021).

**Figure 7.**
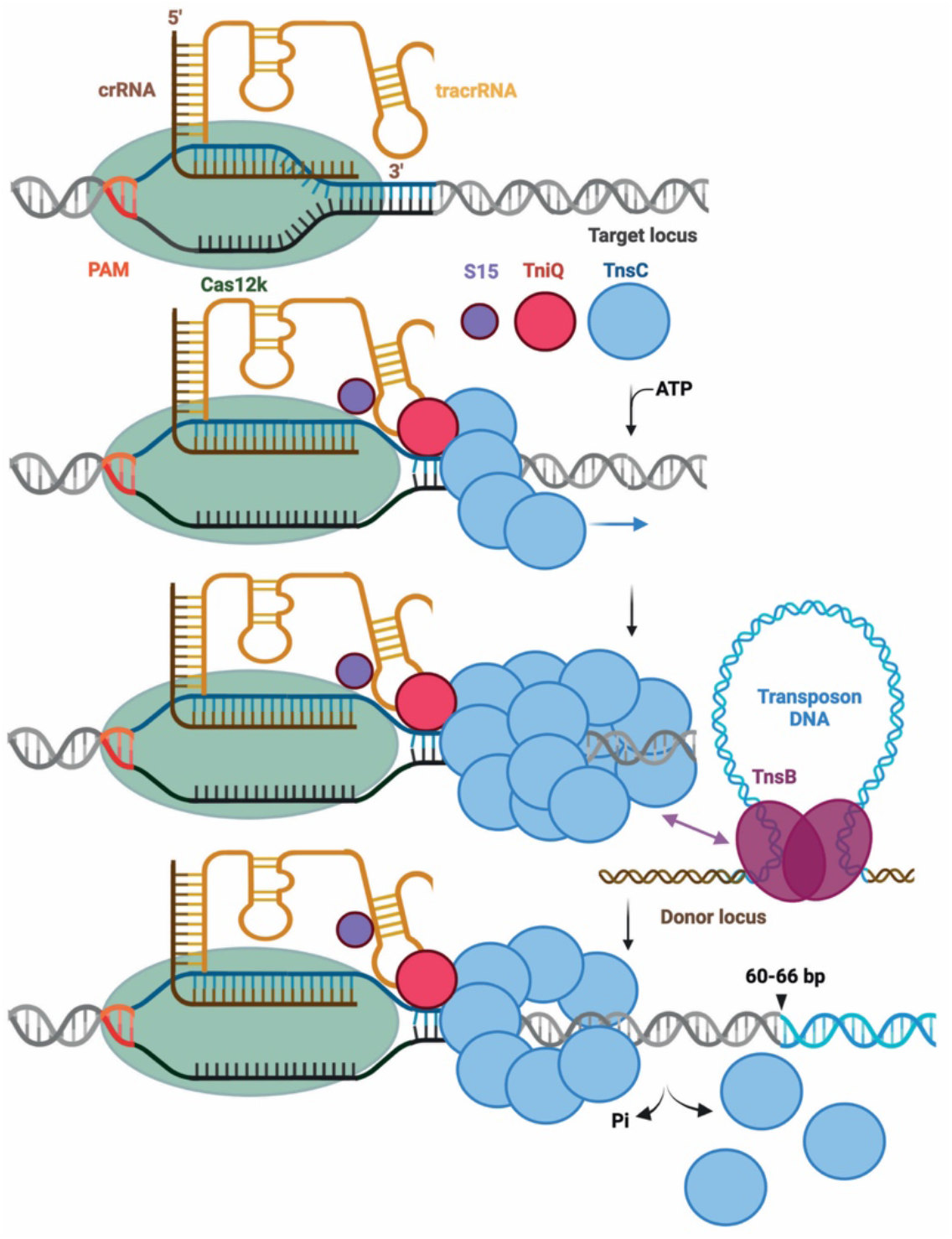
Mechanism of RNA-mediated assembly in type V CRISPR-associated transposons. Mechanistic model for the recruitment of the transposition machinery by the RNA-guided Cas12k complex in type V-K CRISPR-associated transposons. Cas12k in association with a crRNA-tracrRNA dual guide RNA initally binds target DNA to form a partial R-loop structure. Full R-loop formation occurs upon recruitment of S15, TniQ and TnsC. TniQ recognizes specific regions of the tracrRNA and primes oligomerization of TnsC filaments by bridging the first two TnsC protomers. The ribosomal protein S15 assists facilitates productive complex assembly by interacting with the tracrRNA and Cas12k. The resulting TnsC oligomer provides a recruitment platform for TnsB, which triggers TnsC depolymerization to expose the insertion site and catalyzes transposon DNA insertion.

Overall, the transposon recruitment mechanism of type V CRISPR-associated transposons is thus likely to be analogous to that of type I CRISPR-associated transposons in that they both rely on TniQ-dependent assembly of TnsC. However, in type I systems TniQ is an integral component of the Cascade complex (Halpin-Healy et al., 2019; Jia et al., 2020; Li et al., 2020). In type V, TniQ does not form a stable complex on its own with Cas12k-sgRNA and is instead recruited cooperatively together with TnsC (and S15, as discussed below). It is nevertheless possible that TniQ may also interact with TnsC in a Cas12k-independent manner or with other biological R-loop structures, which might lead to off-target transposon recruitment and thus explain why type V CRISPR-transposon systems appear to be less specific than type I systems and more prone to off-target transposon insertion (Strecker et al., 2019; Vo et al., 2021a).

In view of the transposon recruitment complex structure, the previously characterized distance between the Cas12k target site and the transposon insertion site (60-66 bp downstream from the PAM) implies that formation of four complete helical turns of the TnsC filament is required for TnsB recruitment and transposon insertion. As TnsC assembles into hexameric rings on dsDNA in the presence of ADP (Park et al., 2021), it is thought that TnsB-stimulated disassembly of TnsC filaments would result in a TnsC hexamer remaining bound to the Cas12k-TniQ R-loop complex, and the physical footprint of the resulting complex could explain the distance requirement for insertion site selection. However, modeling the structure of this complex by superposing an ADP-bound TnsC hexamer onto the Cas12k-TnsC transposon recruitment complex suggests that the insertion site would be located approximately 28-34 bp from the edge of the TnsC hexamer. It is possible that the discrepancy might be due to the large physical footprint of the TnsB-transposon complex, as indicated by a recent structural study of the *E. coli* Tn7 TnsB bound to transposon end DNA (Kaczmarska et al., 2022). The molecular ruler mechanism determining the distance between the Cas12k target and transposon insertion sites thus remains unclear and awaits further investigation. Nevertheless, the structure of the Cas12k-transposon recruitment complex establishes a paradigm for type I CRISPR-transposon systems and other Tn7-like transposons that rely on the interaction between a sequence-specific DNA binding protein and an oligomeric AAA+ ATPase for site-specific transposon insertion (Choi et al., 2014; Mitra et al., 2010; Shen et al., 2022).

Finally, our structural and functional data indicate that the bacterial ribosomal protein S15 is an integral component of the type V CRISPR-transposon system, as it allosterically stimulates the assembly of TniQ and TnsC in the Cas12k-transposon recruitment complex, thereby enhancing RNA-guided transposition *in vitro*. The involvement of a host-encoded “housekeeping” factor in the activity of a CRISPR effector complex is so far unprecedented. These findings have important implications for the genome engineering application of CRISPR-associated transposons. Type V CAST systems have not yet been demonstrated to support RNA-guided transposition in eukaryotic cells and it is conceivable that this is in part because our description of this systems has hitherto been incomplete. In sum, this work sheds light on a fundamental step in the biological mechanism of CRISPR-associated transposon systems and lays the mechanistic foundation for their development as next-generation genome engineering technologies.

## Supplemental Figures

**Figure S1.**
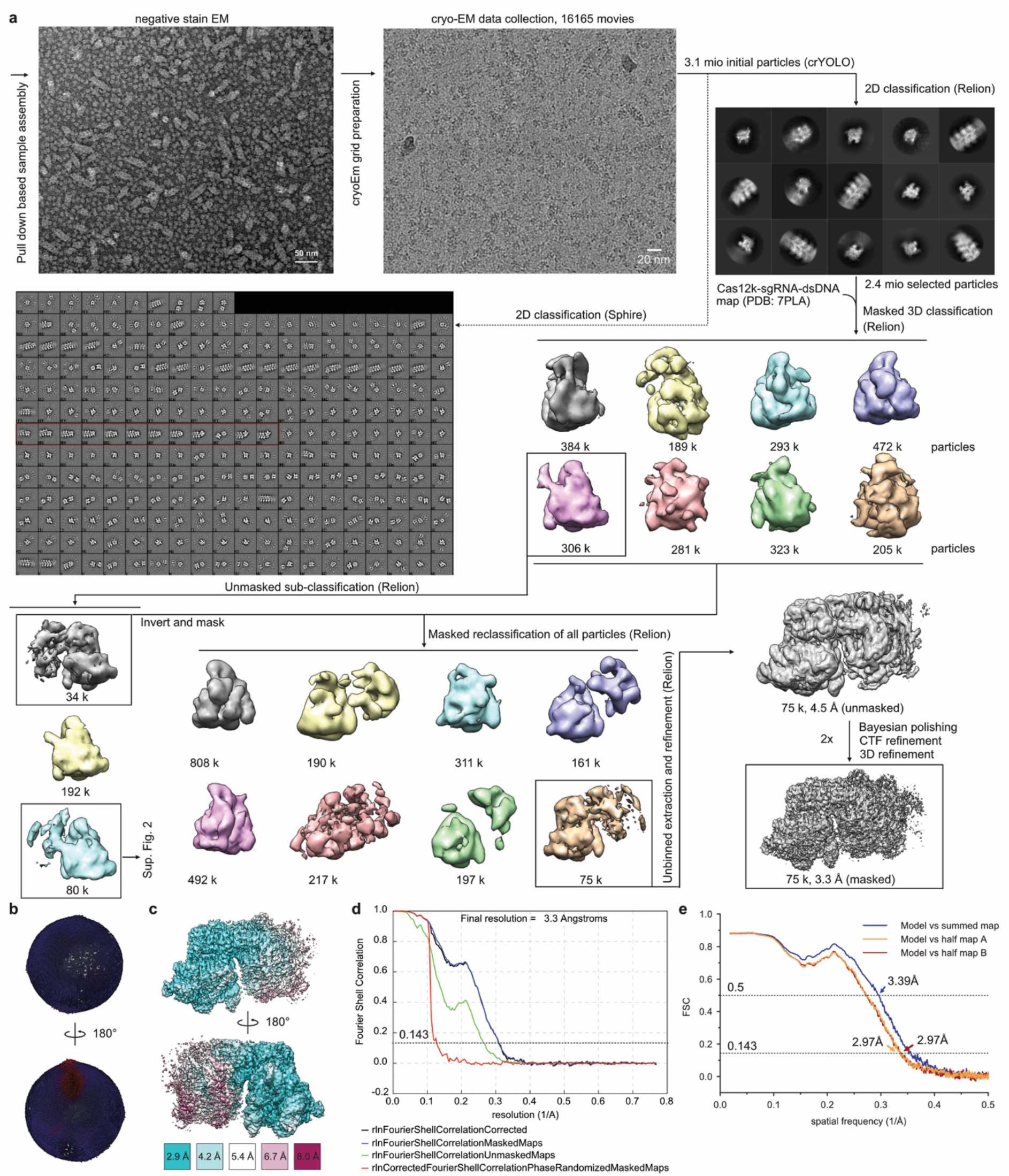
Cryo-EM data processing workflow for the Cas12k-TsnC transposon recruitment complex. (**A**) Cryo-EM image processing workflow for the Cas12k-transposon recruitment complex. Representative negative stain EM and cryo-EM micrographs are shown at 98,000x and 130,000x magnifications, respectively. (**B**) Angular distribution plotted on the density map. (**C**) Final cryo-EM density map colored according to local resolution. (**D**) Fourier Shell Correlations (FSC) of the reconstruction from two independently refined half-maps. The gold-standard cutoff (FSC = 0.143) is marked with a black dotted line. (**E**) Validation of Cas12k-transposon recruitment complex structure model.

**Figure S2.**
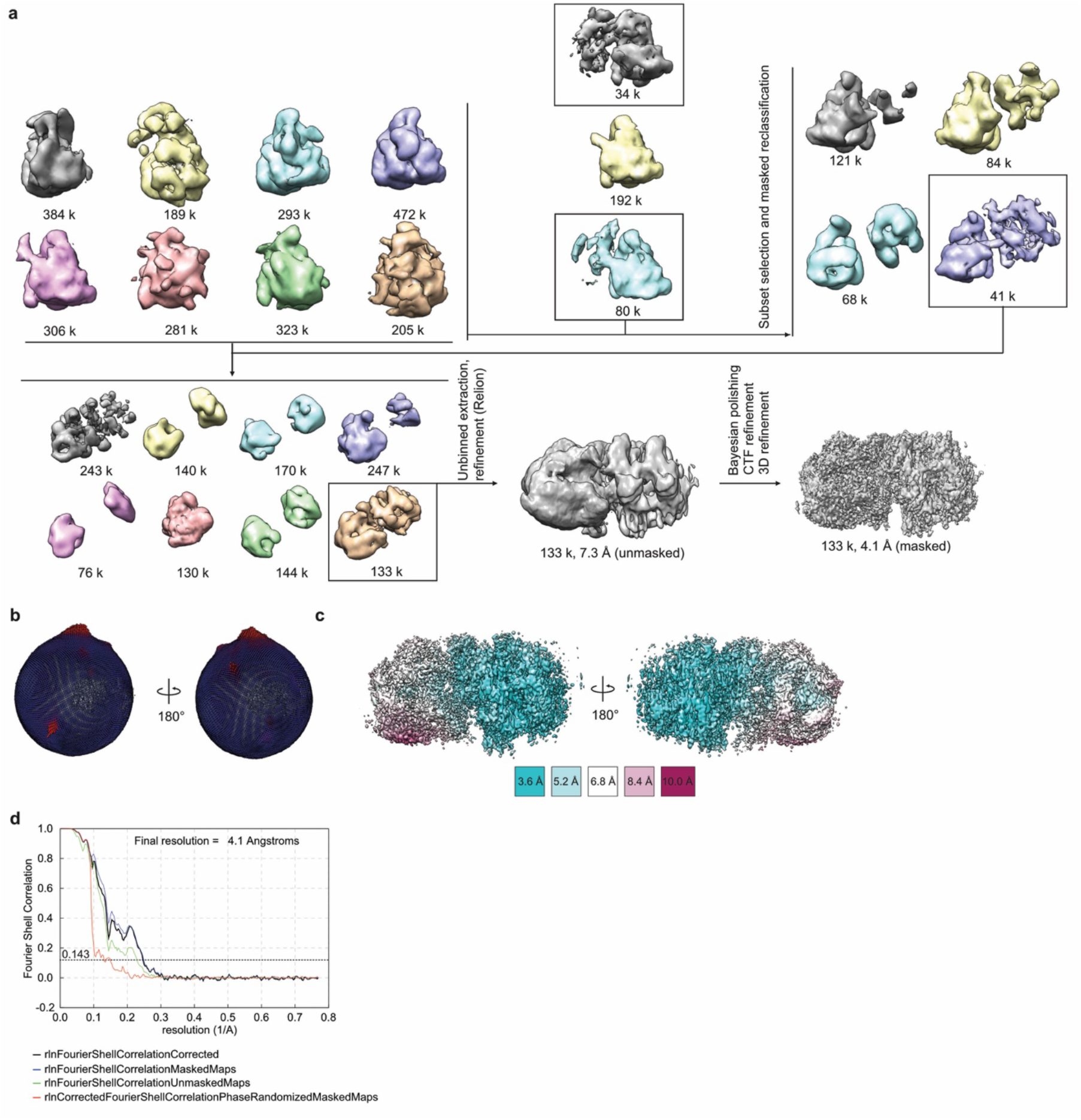
Cryo-EM data processing workflow for the Cas12k-TnsC non-productive complex. (A) Continued details of the cryo-EM image processing workflow from Fig. S1. (**B**) Angular distribution plotted on the resulting density map. (**C**) Final electron density map colored according to the local resolution. (**D**) Fourier Shell Correlations (FSC) of the reconstruction from two independently refined half-maps.

**Figure S3.**
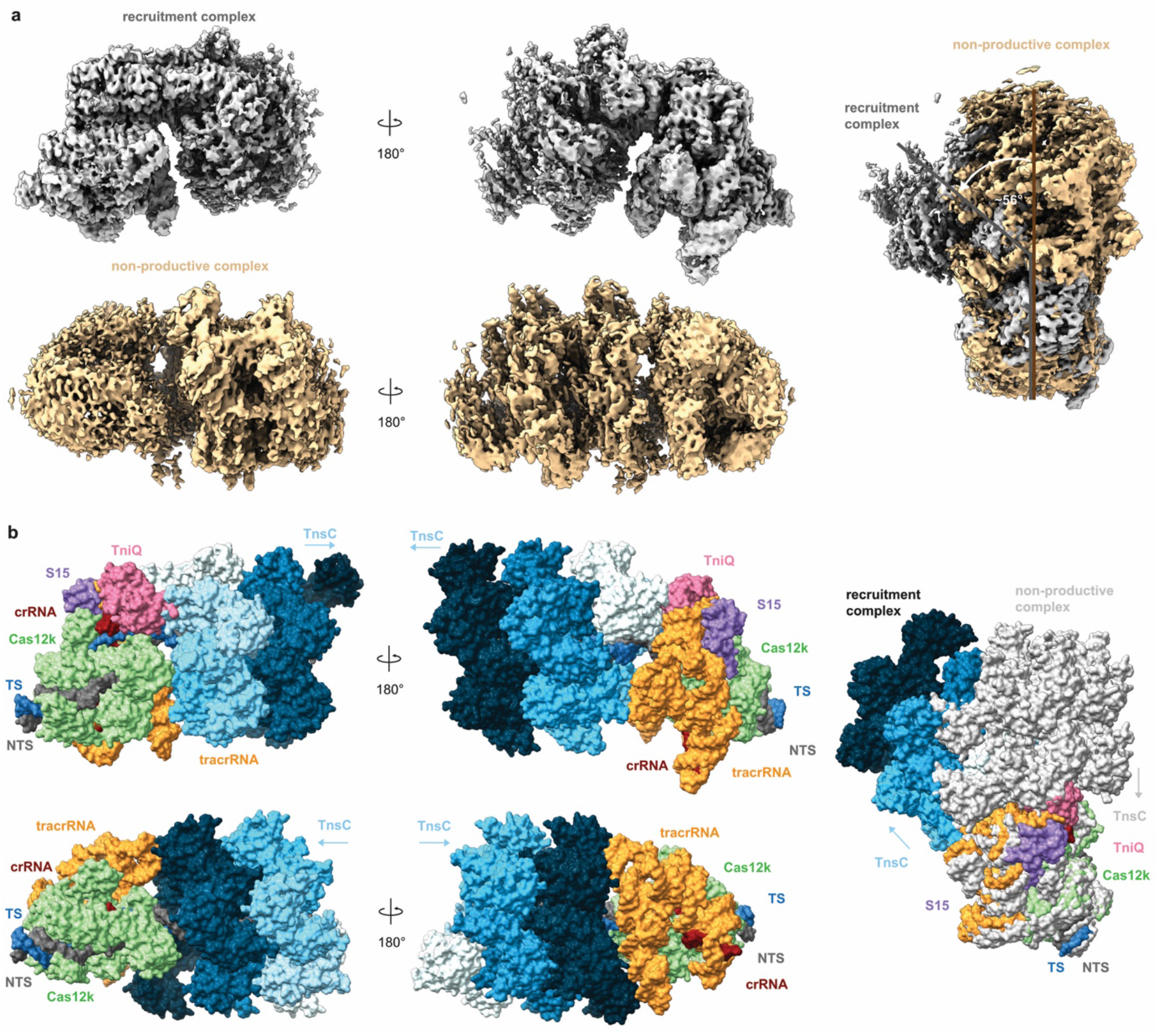
Structural comparison between the Cas12k-transposon recruitment complex and the Cas12k-TnsC non-productive complex. (**A**) Cryo-EM density maps of the Cas12k-transposon recruitment complex (top) and the Cas12k-TnsC non-productive complex (bottom). Side views and structural superpositions are shown. Proteins are shown in surface representation. The DNA in the transposon recruitment complex is bent by ∼56° relative to the non-productive complex. (**B**) Atomic models, shown in surface representation, of the Cas12k-transposon recruitment complex (top) and the Cas12k-TnsC non-productive complex (bottom).

**Figure S4.**
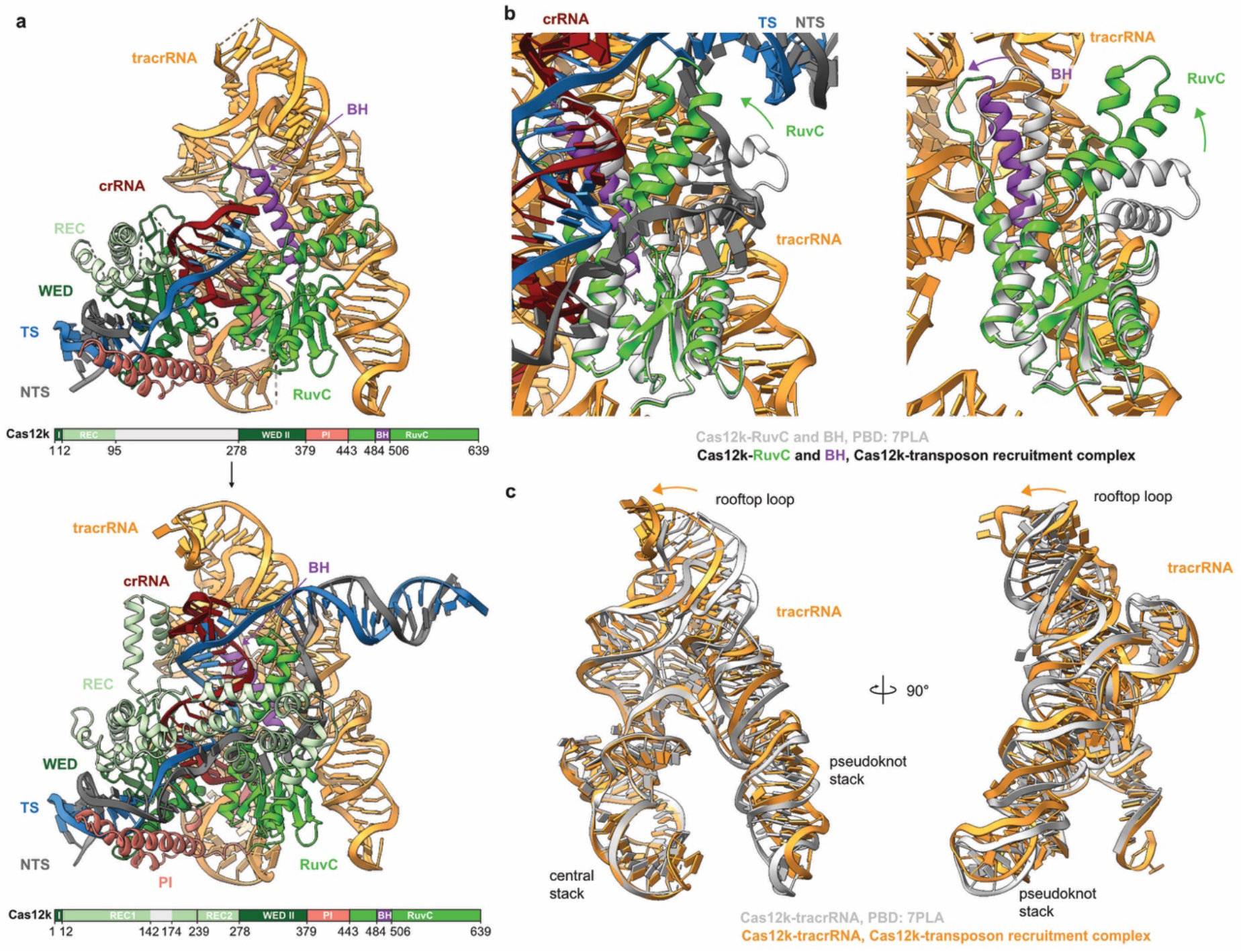
Structural rearrangements in Cas12k and guide RNA upon R-loop completion. (**A**) Structural models of the Cas12k-sgRNA-target DNA complex (top) and the Cas12k-transposon recruitment complex (bottom), shown in the same orientation. Domain architectrue of Cas12k is shown below each model. REC, recognition lobe. WED, wedge domain. PI, PAM interacting domain. BH, bridging helix. TS, target DNA strand; NTS, non-target DNA strand. (**B**) Structural superpositions of the RuvC and BH domains in the Cas21k-sgRNA-target DNA and Cas12k-transposon recruitment complexes. (**C**) Structural superposition of the tracrRNA part of the sgRNA in the Cas12k-sgRNA-target DNA (PDB ID:7PLA, shown in grey) and the Cas12k-transposon recruitment complex (orange).

**Figure S5.**
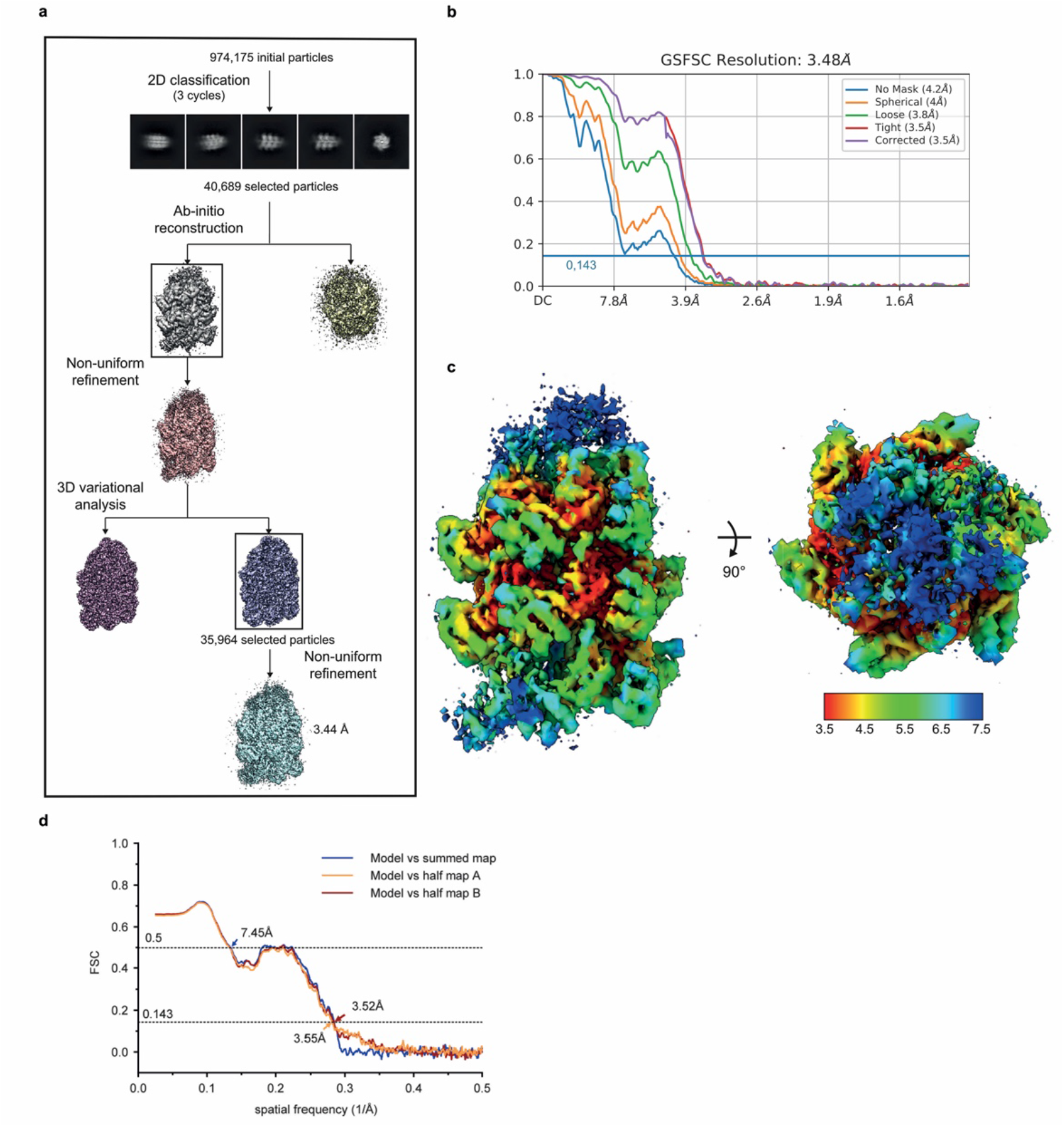
Cryo-EM analysis of TniQ-capped TnsC filament. (**A**) Cryo-EM image processing workflow for the TniQ-capped TnsC filament complex. (**B**) Fourier Shell Correlations (FSC) of ShTnsC-DNA-ShTniQ reconstruction from two independently refined half-maps. The gold-standard cutoff (FSC = 0.143) is marked with a blue line. (**C**) Final electron density map colored according to the local resolution. (**D**) Fourier Shell Correlations (FSC) of the reconstruction from two independently refined half-maps.

**Figure S6.**
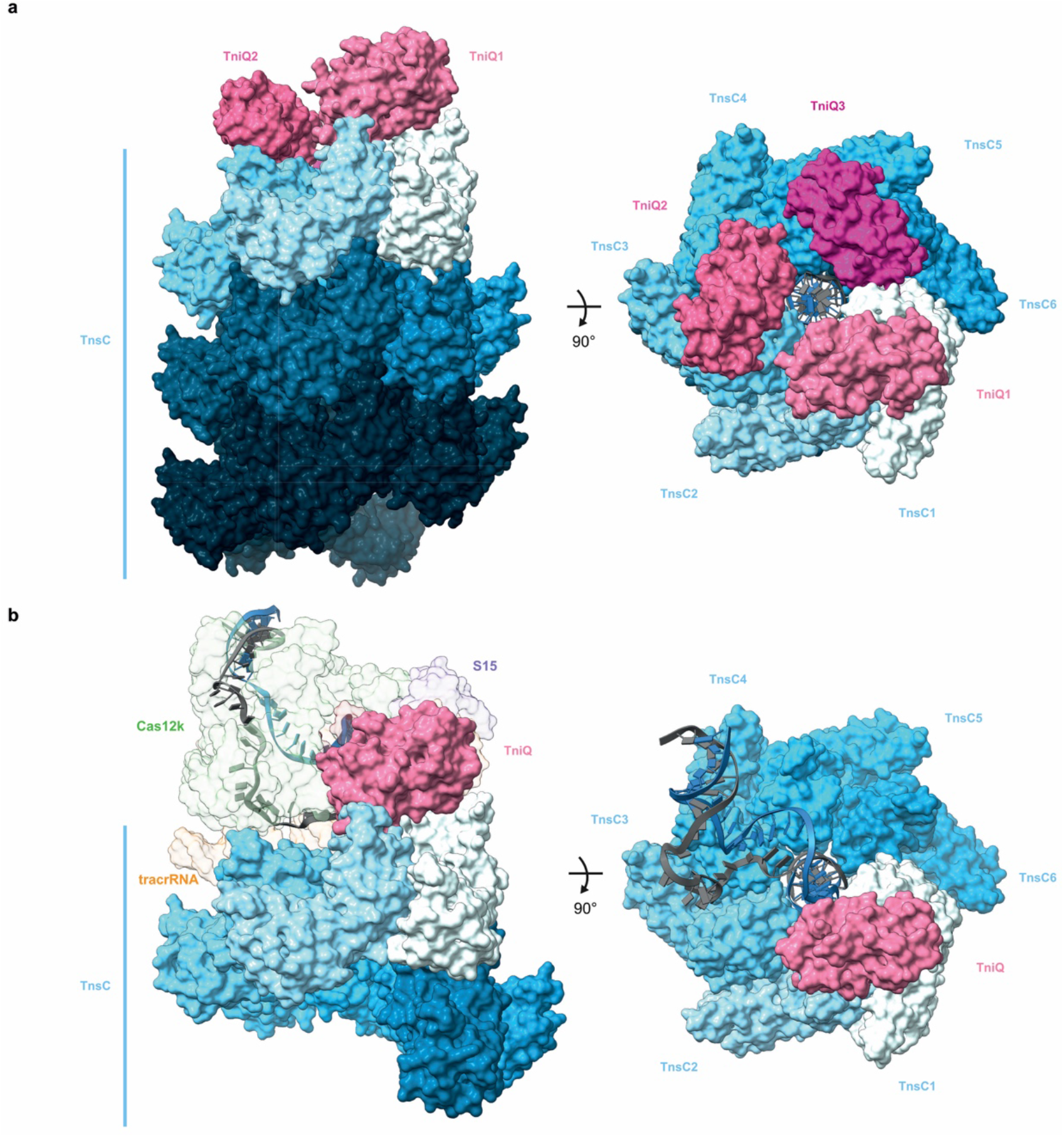
Structural comparisons of TniQ-capped TnsC filament and Cas12k-transposon recruitment complex. (**A**)Side and top views of the TniQ-capped TnsC filament. Proteins are shown in surface representation. (**B**)Side and top views of the Cas12k-transposon recruitment complex, with TniQ shown in the same orientation as TniQ1 in (A). In the top view Cas12k, S15 and tracrRNA are hidden to visualize the contacts between TniQ and TnsC.

**Figure S7.**
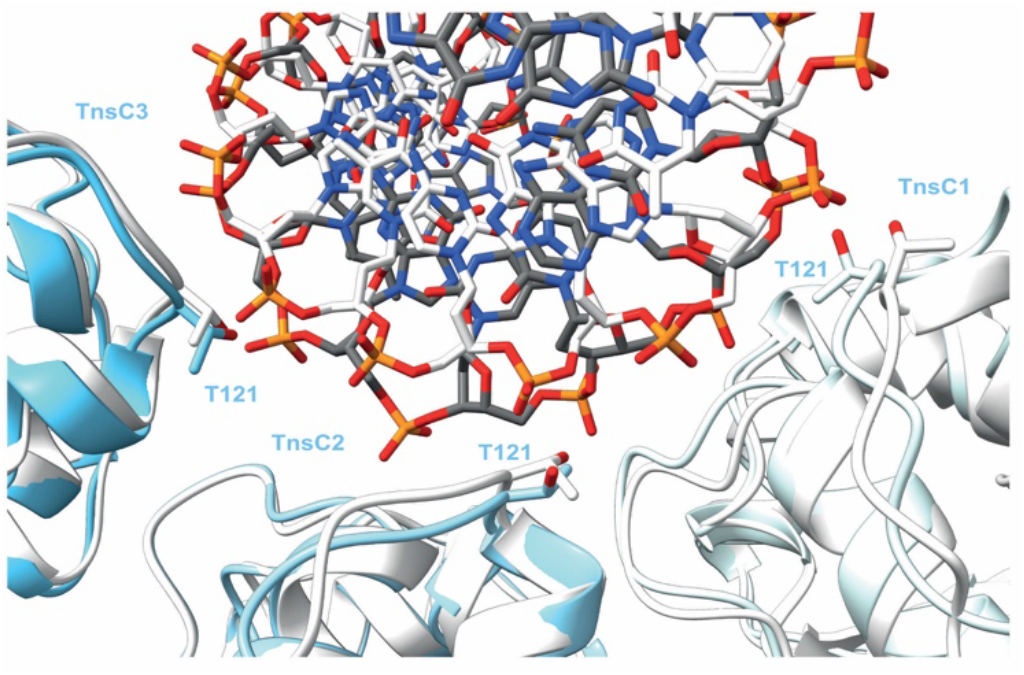
Comparison of DNA recognition by the Cas12k-transposon recruitment complex and the TniQ-capped TnsC filament. Structural overlay of three consecutive TnsC protomers (TnsC1-TnsC3) in the Cas12k-transposon recruitment complex (colored protomers, gray DNA) and in the TniQ-capped TnsC filament (white protomers and DNA) and the associated DNA (shown in stick representation).

**Figure S8.**
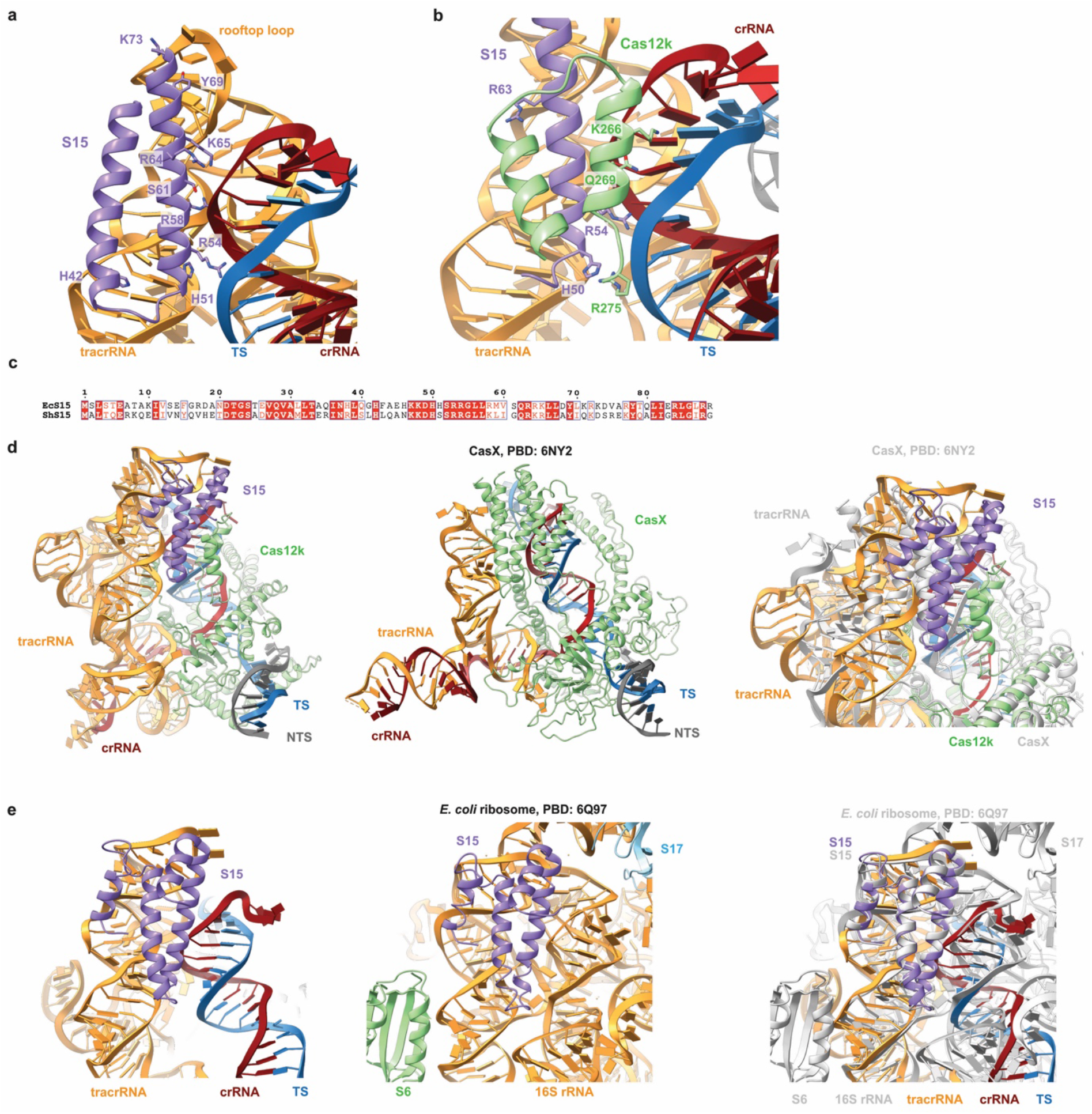
Interactions and conservation of the ribosomal protein S15. (**A**) Zoomed-in view of S15 interactions with the tracrRNA and crRNA:TS-DNA duplex. (**B**) Zoomed-in view of S15 contacts with the Cas12k REC2 domain. (**C**) Sequence alignment of the ribosomal proteins S15 from *E. coli* (EcS15) and *S. hoffmanni* (ShS15). (**D**) Structural models of the Cas12k-transposon recruitment complex (left), CasX-sgRNA-target DNA complex (middle, PDB: 6NY2) and a superposition of both focused on S15 (right). (**E**) Zoomed-in views of EcS15 interactions with tracrRNA in the Cas12k-transposon recruitment complex (left), 16S rRNA in the *E. coli* ribosome (middle, PDB ID: 6Q97 (Rae et al., 2019)) and a superposition of both focused on S15 (right).

## Supplemental Tables

**Table S1.** Cryo-EM data collection, refinement and validation statistics for the ShCas12k-sgRNA-dsDNA-S15-TniQ-TnsC complex structure and the TnsC-ATP-dsDNA-TniQ structure.

|  | Cas12k-sgRNA-dsDNA-S15-TniQ-TnsC<br>(EMDB- xxxx)<br>(PDB xxx) | TnsC-ATP-dsDNA-TniQ<br>(EMDB- yyyy)<br>(PDB yyy) |
| --- | --- | --- |
| **Data collection and processing** |  |  |
| Magnification | 130,000 | 130,000 |
| Voltage (kV) | 300 | 300 |
| Electron exposure (e-/Å<sup>2</sup>) | 67.68 | 66.036 |
| Defocus range (µm) | -1.0 to -2.4 (0.2 steps) | -1.0 to -2.4 (0.2 steps) |
| Pixel size (Å) | 0.325 | 0.325 |
| Symmetry imposed | C1 | C1 |
| Initial particle images (no.) | 16,165 | 10,436 |
| Final particle images (no.) | 15,296 | 10,176 |
| Map resolution (Å) | 3.28 | 3.48 |
| FSC threshold | 0.143 | 0.143 |
| Map resolution range (Å) | 2.9-8.0 | 3.4-7.5 |
| **Refinement** |  |  |
| Initial model used (PDB code) | 7PLA & 7PLH | 7OXD & 7PLH |
| Model resolution (Å) | 3.5 | 3.5 |
| FSC threshold | 0.5 | 0.143 |
| Map sharpening *B* factor (Å<sup>2</sup>) | -66.35 | -47.3 |
| Model composition |  |  |
| Non-hydrogen atoms | 28381 | 18984 |
| Protein residues | 2669 | 2285 |
| Nucleotide residues | 328 | 32 |
| Ligands | Zn : 2 MG :7 ATP :7 | Zn : 6 MG :7 ATP :7 |
| *B* factors (Å<sup>2</sup>) min/max/mean |  |  |
| Protein | 0.00/449.71/194.31 | 30.00/428.64/201.00 |
| Nucleotide | 17.06/522.32/138.45 | 30.00/30.00/30.00 |
| Ligand | 126.42/309.10/243.23 | 30.00/299.56/234.94 |
| R.m.s. deviations |  |  |
| Bond lengths (Å) | 0.008 | 0.005 |
| Bond angles (°) | 0.692 | 1.054 |
| Validation |  |  |
| MolProbity score | 1.84 | 2.05 |
| Clashscore | 16.21 | 30.40 |
| Poor rotamers (%) | 0.18 | 0.64 |
| Ramachandran plot |  |  |
| Favored (%) | 97.36 | 97.62 |
| Allowed (%) | 2.64 | 2.38 |
| Disallowed (%) | 0.00 | 0.00 |
| Ramachandran Z score |  |  |
| whole | -0.24 | -0.08 |
| helix | 0.56 | 0.57 |
| sheet | -1.58 | -1.90 |
| loop | -1.09 | -0.68 |

**Table S2** | **Mass spectrometry analysis of TniQ and co-purified proteins**.

**Table S3** | **Oligonucleotides, gBlocks and synthetic genes used in this study**.

## Methods

### Plasmid DNA constructs

The DNA sequences of *Scytonema hofmanni* Cas12k (WP_029636312.1), TnsC (WP_029636336.1), TniQ (WP_029636334.1), TnsB (WP_084763316.1), S15 (WP_029633173.1) and *Escherichia coli* S15 (AP009048.1) proteins were codon optimized for heterologous expression in *Escherichia coli* (*E. coli*) and synthesized by GeneArt (Thermo Fisher Scientific). The ShCas12k gene was inserted into the 1B (Addgene 29653) and 1R (Addgene 29664) plasmids using ligation-independent cloning (LIC), resulting in constructs carrying an N-terminal hexahistidine tag followed by a tobacco etch virus (TEV) protease cleavage site and a N-terminal hexahistidine-StrepII tag followed by a TEV cleavage site, respectively. The ShTnsC gene was inserted into the 1S (Addgene 29659) plasmid to produce a construct carrying a N-terminal hexahistidine and hexahistidine-SUMO tag followed by a TEV cleavage site and the ShTniQ was cloned into the 1C (Addgene 29659) vector generating a construct carrying a hexahistidine-maltose binding protein (6xHis-MBP) tag followed by a TEV cleavage site. The EcS15 and ShS15 genes were inserted into the 1C vector. Point mutations were introduced by Gibson assembly using gBlock gene fragments synthetized by IDT or annealed oligonucleotides provided by Sigma as inserts. The pDonor (Addgene 127921), pHelper (Addgene 127924), and pTarget (Addgene 127926) plasmids (Strecker et al., 2019) used in droplet digital PCR experiments were sourced from Addgene. The PSP1-targeting spacer was cloned into pHelper by Gibson assembly, yielding pHelper-PSP1. Mutations in the sgRNA or in the sequence of the ShCas12k and ShTnsC genes were introduced into the pHelper plasmid by Gibson assembly. Plasmids were cloned and propagated in Mach I cells (Thermo Fisher Scientific) with the exception of pHelper, which was grown in One Shot PIR1 cells (Thermo Fisher Scientific). Plasmids were purified using the GeneJET plasmid miniprep kit (Thermo Fisher Scientific) and verified by Sanger sequencing. All sequence of primers, gBlocks and oligonucleotides used for cloning are provided in **Table S3**.

### Protein expression and purification

For expression of ShCas12k constructs, hexahistidine-Strep II-tagged and hexahistidine-tagged ShCas12k proteins were expressed in *E. coli* BL21 Star (DE3) cells. Cell cultures were grown at 37 °C shaking at 100 rpm until reaching an OD_600_ of 0.6 and protein expression was induced with 0.4 mM IPTG (isopropyl-β-d-thiogalactopyranoside) and continued for 16 h at 18 °C. Harvested cells were resuspended in 20 mM Tris-HCl pH 8.0, 500 mM NaCl, 5 mM Imidazole, 1 μg/mL Pepstatin, 200 μg/mL AEBSF and lysed in a Maximator Cell homogenizer at 2,000 bar and 4 °C. The lysate was cleared by centrifugation at 40,000 x *g* for 30 min at 4 °C and applied to two 5 mL Ni-NTA cartridges connected in tandem. The column was washed with 100 mL of 20 mM Tris-HCl pH 8.0, 500 mM NaCl, 5 mM Imidazole before elution with 50 mL of 20 mM Tris-HCl pH 8.0, 500 mM NaCl, 250 mM Imidazole. Elution fractions were pooled and dialyzed overnight against 20 mM HEPES-KOH pH 7.5, 250 mM KCl, 1 mM EDTA, 1 mM DTT in the presence of tobacco etch virus (TEV) protease. Dialyzed proteins were loaded onto a 5 mL HiTrap Heparin HP column (GE Healthcare) and eluted with a linear gradient of 20 mM HEPES-KOH pH 7.5, 1 M KCl. Elution fractions were pooled, concentrated using 30,000 molecular weight cut-off centrifugal filters (Merck Millipore) and further purified by size exclusion chromatography using a Superdex 200 (16/600) column (GE Healthcare) in 20 mM HEPES-KOH pH 7.5, 250 mM KCl, 1 mM DTT yielding pure, monodisperse proteins. Purified proteins were concentrated to 10-15 mg mL^−1^, flash frozen in liquid nitrogen and stored at -80 °C until further usage.

Expression of wild-type ShTnsC and ShTniQ was performed in *E. coli* BL21 Rosetta2 (DE3) cells. Cells were grown in LB medium until reaching an OD_600_ of 0.6 and expression was induced by addition of 0.4 mM IPTG. Proteins were expressed at 18 °C for 16 h. For ShTnsC, the cells were harvested and resuspended in lysis buffer containing 20 mM Tris-HCl pH 7.5, 500 mM NaCl, 5% glycerol and 10 mM imidazole supplemented with EDTA-free protease inhibitor (Roche). The cell suspension was lysed by ultrasonication and the lysate was cleared by centrifugation at 40,000 x *g* for 40 min. Cleared lysate was applied to a 5 mL Ni-NTA cartridge (Qiagen). The column was washed in two steps with lysis buffer supplemented with 25 mM and 100 mM imidazole, and bound proteins were eluted with 25 mL of same buffer supplemented with 500 mM imidazole pH 7.5. Eluted proteins were dialysed overnight against 20 mM Tris-HCl pH 7.5, 250 mM NaCl, 5% glycerol, 1 mM DTT in the presence of TEV protease. The protein was further purified using a 5 mL HiTrap HP Heparin column (GE Healthcare) and eluted with a buffer containing 20 mM Tris-HCl pH 7.5, 700 mM NaCl, 5% glycerol and 1 mM DTT. The eluted fraction was concentrated and further purified by size exclusion chromatography using an S200 increase (10/300 GL) column (GE Healthcare) in 20 mM Tris-HCl pH 7.5, 500 mM NaCl, 1 mM DTT, yielding pure, monodisperse proteins. Purified ShTnsC was concentrated to 1-2 mg mL^−1^ using 30,000 kDa molecular weight cut-off centrifugal filters (Merck Millipore) and flash-frozen in liquid nitrogen.

For ShTnsB, cells were harvested and resuspended in lysis buffer containing 20 mM Tris-HCl pH 8.0, 500 mM NaCl, 5% glycerol and 5 mM Imidazole supplemented with EDTA-free protease inhibitor (Roche) and lysed by ultrasonication. Lysate was clarified by centrifugation at 40,000 x *g* for 40 min. The lysate was applied to a 5 mL Ni-NTA cartridge (Qiagen) and the column was washed with lysis buffer supplemented with 25 mM imidazole. The protein was eluted with 20 mM Tris-HCl pH 7.5, 500 mM NaCl, 5% glycerol and 200 mM Imidazole. Eluted proteins were dialysed overnight against 20 mM Tris-HCl pH 8.0, 200 mM NaCl, 1 mM DTT in the presence of TEV protease and subsequently purified using a 5 mL HiTrap HP Heparin column (GE Healthcare), eluting with a buffer containing 20 mM Tris-HCl pH 8.0, 400 mM NaCl, and 1 mM DTT. The eluted fraction was concentrated and further purified by size exclusion chromatography using a Superdex (16/600) column (GE Healthcare) in 20 mM Tris-HCl pH 8.0, 400 mM NaCl, and 1 mM DTT. Purified ShTnsB was concentrated to 7 mg mL^−1^, and flash-frozen in liquid nitrogen. For the ShTnsB G190-F584 and ShTnsB G190-L494 constructs, cells were harvested and resuspended in 20 mM Tris-HCl pH 7.5, 500 mM NaCl, 5% glycerol and 5 mM imidazole supplemented with EDTA-free protease inhibitor (Roche) and lysed by ultrasonication. Cleared lysate was applied to a 5 mL Ni-NTA cartridge (Qiagen) and the column was washed in two steps with lysis buffer supplemented with 25 and 50 mM imidazole. The proteins were eluted with 20 mM Tris-HCl pH 7.5, 500 mM NaCl, 5% glycerol and 125 mM imidazole and dialysed overnight against 20 mM Tris-HCl pH 7.5, 500 mM NaCl, 5% glycerol and 4 mM beta-mercaptoethanol in the presence of TEV. Dialysed proteins were passed through a 5 mL Ni-NTA cartridge and washed with buffer containing 20 mM Tris-HCl pH 7.5, 500 mM NaCl, 5% glycerol and 25 mM imidazole, and further purified by size exclusion chromatography using an Superdex (16/600) column (GE Healthcare) in 20 mM Tris-HCl pH 7.5, 250 mM NaCl, 1 mM DTT. Purified ShTnsB proteins were concentrated to 13–24 mg mL^−1^ and flash-frozen in liquid nitrogen.

For ShTniQ, cells were harvested and resuspended in 20 mM Tris-HCl pH 7.8, 500 mM NaCl, 5% glycerol and 5 mM imidazole supplemented with EDTA-free protease inhibitor (Roche) and lysed by ultrasonication. The cleared lysate was applied to a 5 mL Ni-NTA cartridge (Qiagen) and the column was washed in two steps with lysis buffer supplemented with 25 and 50 mM imidazole. The protein was eluted with lysis buffer supplemented with 300 mM imidazole. Eluted protein was dialysed overnight against 20 mM Tris-HCl pH 7.8, 500 mM NaCl, 1 mM DTT in the presence of TEV protease. Dialysed protein was passed through a 5 mL MBPTrap column (GE Healthcare). The flow-through fraction was concentrated and further purified by size exclusion chromatography using a Superdex 200 (16/600) column (GE Healthcare) in 20 mM Tris-HCl pH 7.8, 250 mM NaCl, 1 mM DTT. Purified ShTniQ was concentrated to 10 mg mL^−1^ using 10,000 kDa molecular weight cut-off centrifugal filters (Merck Millipore) and flash-frozen in liquid nitrogen. To produce recombinant ShTniQ protein free of the *E. coli* S15 contaminant, the protocol was adjusted as follows. Cells were harvested and resuspended in 20 mM Tris-HCl pH 7.5, 500 mM NaCl, 5% glycerol and 5 mM imidazole supplemented with EDTA-free protease inhibitor (Roche) and lysed by ultrasonication. The lysate was cleared by centrifugation at 40,000 x *g* for 30 min at 4 °C and applied to two 5 mL Ni-NTA cartridges connected in tandem. The column was washed with 150 mL of 20 mM Tris-HCl pH 7.5, 500 mM NaCl, 5 mM imidazole, 5 % glycerol before elution with 50 mL of 20 mM Tris-HCl pH 7.5, 500 mM NaCl, 500 mM imidazole, 5 % glycerol into two 5 mL MBP-trap cartridges (GE Healthcare) connected in tandem before removal of the Ni-NTA cartridges. The MBP-trap column was washed with 50 mL of 20 mM Tris-HCl pH 7.5, 500 mM NaCl, 500 mM imidazole, 5 % glycerol before elution with 50 mL of 20 mM Tris-HCl pH 8.0, 500 mM NaCl, 500 mM Imidazole, 5 % glycerol, 10 mM maltose. Elution fractions were pooled and dialyzed overnight against 20 mM HEPES-KOH pH 7.5, 100 mM KCl, 1 mM DTT in the presence of tobacco etch virus (TEV) protease. Dialyzed proteins were loaded onto two 5 mL HiTrap Heparin HP columns (GE Healthcare) connected in tandem, and eluted with a linear gradient of 20 mM HEPES-KOH pH 7.5, 1 M KCl. Elution fractions were pooled, concentrated using 3,000 molecular weight cut-off centrifugal filters (Merck Millipore) and further purified by size exclusion chromatography using a Superdex 75 (16/600) column (GE Healthcare) in 20 mM Tris-HCl pH 7.5, 250 mM NaCl, 1 mM DTT yielding pure, monodisperse proteins. Purified proteins were concentrated to 1.5-5.0 mg mL^−1^, flash frozen in liquid nitrogen and stored at -80 °C until further usage. Absence of the *E. coli* S15 in the purified ShTniQ produced using the revised protocol was confirmed by mass spectrometry as described below (see also **Table S2**).

For expression of EcS15 and ShS15, hexahistidine-MBP-tagged proteins were expressed in *E. coli* BL21 Rosetta2 (DE3) cells. Cell cultures were grown at 37 °C shaking at 100 rpm until reaching an OD_600_ of 0.6 and protein expression was induced with 0.4 mM IPTG (isopropyl-β-d-thiogalactopyranoside) and continued for 16 h at 18 °C. Harvested cells were resuspended in 20 mM Tris-HCl pH 8.0, 500 mM NaCl, 5 mM Imidazole, 1 μg/mL Pepstatin, 200 μg/mL AEBSF and lysed by ultrasonication at 4 °C. The lysate was cleared by centrifugation at 40,000 x *g* for 30 min at 4 °C and applied to two 5 mL Ni-NTA cartridges connected in tandem. The column was washed with 100 mL of 20 mM Tris-HCl pH 8.0, 500 mM NaCl, 5 mM Imidazole before elution with 50 mL of 20 mM Tris-HCl pH 8.0, 500 mM NaCl, 250 mM Imidazole. Elution fractions were pooled and dialyzed overnight against 20 mM HEPES-KOH pH 7.5, 150 mM KCl, 1 mM DTT in the presence of tobacco etch virus (TEV) protease. Dialyzed proteins were loaded onto a 5 mL HiTrap Heparin HP column (GE Healthcare) and eluted with a linear gradient of 20 mM HEPES-KOH pH 7.5, 1 M KCl. Elution fractions were pooled, concentrated using 3,000 molecular weight cut-off centrifugal filters (Merck Millipore) and further purified by size exclusion chromatography using a Superdex 200 (16/600) column (GE Healthcare) in 20 mM HEPES-KOH pH 7.5, 250 mM KCl, 1 mM DTT yielding pure, monodisperse proteins. Purified proteins were concentrated to 2-5 mg mL^−1^, flash frozen in liquid nitrogen and stored at -80 °C until further usage.

### Mass spectrometry analysis

Samples resulting from purification of TniQ following the Protocol 1 were treated as follows prior to digestion. Proteins were precipitated with trichloroacetic acid (TCA; Sigma-Aldrich) at a final concentration of 5% and washed twice with ice-cold acetone. The protein pellet was resuspended in 45 μL of digestion buffer (10 mM Tris, 2 mM CaCl2, pH 8.2). Samples resulting from purification of TniQ following the Protocol 2 were directly diluted in digestion buffer (10 mM Tris, 2 mM CaCl2, pH 8.2) and reduced with 2 mM TCEP(tris(2-carboxyethyl)phosphine) and alkylated with 15 mM chloroacetamide at 30°C for 30 min. Digestion was performed for all samples using the same procedure: 500 ng of Sequencing Grade Trypsin (Promega) were added for digestion carried out in a microwave instrument (Discover System, CEM) for 30 min at 5 W and 60 °C. The samples were dried to completeness and re-solubilized in 20 μL of MS sample buffer (3% acetonitrile, 0.1% formic acid).

LC-MS/MS analysis was performed on an Q Exactive HF mass spectrometer (Thermo Scientific) equipped with a Digital PicoView source (New Objective) and coupled to an M-Class UPLC (Waters). Solvent composition at the two channels was 0.1% formic acid for channel A and 0.1% formic acid, 99.9% acetonitrile for channel B. For each sample, 80 ng of peptides were loaded on a commercial ACQUITY UPLC M-Class Symmetry C18 Trap Column (100Å, 5 μm, 180 μm x 20 mm, Waters) connected to a ACQUITY UPLC M-Class HSS T3 Column (100Å, 1.8 μm, 75 μm X 250 mm, Waters). The peptides were eluted at a flow rate of 300 nL/min. After a 3 min initial hold at 5% B, a gradient from 5 to 35 % B in 60 min and 35 to 40% B in additional 5 min was applied. The column was cleaned after the run by increasing to 95 % B and holding 95 % B for 10 min prior to re-establishing loading condition. The mass spectrometer was operated in data-dependent mode (DDA), funnel RF level at 60 %, and heated capillary temperature at 275 °C. Full-scan MS spectra (350−1500 m/z) were acquired at a resolution of 120’000 for sample from Protocol 2 and of 70’000 for sample from Protocol 1 at 200 m/z after accumulation to a target value of 3’000’000, followed by HCD (higher-energy collision dissociation) fragmentation on the twelve most intense signals per cycle. Ions were isolated with a 1.2 m/z isolation window and fragmented by higher-energy collisional dissociation (HCD) using a normalized collision energy of 28% for sample from Protocol 2 and of 25% for sample from Protocol 1. HCD spectra were acquired at a resolution of 30’000 and a maximum injection time of 50 ms for sample from Protocol 2 and at a resolution of 35’000 and a maximum injection time of 120 ms for sample from Protocol 1. The automatic gain control (AGC) was set to 100’000 ions. Charge state screening was enabled and singly and unassigned charge states were rejected. Precursor masses previously selected for MS/MS measurement were excluded from further selection for 30 s, and the exclusion window tolerance was set at 10 ppm. The samples were acquired using internal lock mass calibration on m/z 371.1010 and 445.1200. The mass spectrometry proteomics data were handled using the local laboratory information management system (LIMS) (Turker et al., 2011).

The acquired raw MS data were processed by MaxQuant (version 1.6.2.3), followed by protein identification using the integrated Andromeda search engine (Cox and Mann, 2008). Spectra were searched against a Uniprot E. coli proteome database (UP000000625, taxonomy 83333, version from 2021-07-12), concatenated to its reversed decoyed fasta database and the sequences of the proteins of interest. Acetyl (Protein N-term) oxidation (M) and deamidation (NQ) were set as variable modification. Enzyme specificity was set to trypsin/P allowing a minimal peptide length of 7 amino acids and a maximum of two missed-cleavages. MaxQuant Orbitrap default search settings were used. The maximum false discovery rate (FDR) was set to 0.01 for peptides and 0.05 for proteins. Peptide identifications were accepted if they achieved a false discovery rate (FDR) of less than 0.1% by the Scaffold Local FDR algorithm. Protein identifications were accepted if they achieved an FDR of less than 1.0% and contained at least 2 identified peptides. Data are provided in **Table S2**.

### sgRNA preparation

DNA sequence encoding the T7 RNA polymerase promoter upstream of the ShCas12k-sgRNA was sourced as a gBlock (IDT), cloned into a pUC19 plasmid using restriction digest with BamHI and EcoRI, and confirmed by Sanger sequencing. The sequence encoding the T7 RNA polymerase promoter and sgRNA was amplified by PCR and purified by ethanol precipitation for use as template for *in vitro* transcription with T7 RNA polymerase as described previously (Anders et al., 2015). The transcribed RNA was gel purified, precipitated with 70 % (v/v) ethanol, dried and dissolved in nuclease-free water.

### Cryo-EM sample preparation and data collection: Cas12k-transposon recruitment complex

The complex was generated by stepwise assembly on Strep-tactin matrix as follows. First, the sgRNA was mixed with hexahistidine-StrepII-tagged ShCas12k (Strep-Cas12k) in assembly buffer (20 mM HEPES-KOH pH 7.5, 250 mM KCl, 10 mM MgCl_2_, 1 mM DTT) and incubated 20 min at 37 °C to allow formation of a binary Strep-Cas12k-sgRNA complex. Next, a target dsDNA duplex (**Table S3**) was added to the reaction in a Strep-Cas12k:sgRNA:dsDNA molar ratio of 1:1.2:2 and incubated for 20 min at 37 °C, yielding an assembled Strep-Cas12k-sgRNA-target DNA complex. The final 30 μL reaction contained 20 μg (at a concentration of 8.4 μM, 0.7 mg/mL) Strep-Cas12k, 10.1 μM sgRNA, and 16.9 μM DNA in assembly buffer. The resulting sample was then mixed with 25 μL Strep-Tactin beads (iba) equilibrated in pull-down wash buffer 1 (20 mM HEPES-KOH pH 7.5, 250 mM KCl, 10 mM MgCl_2_, 1 mM DTT, 0.05 % Tween20) and incubated 30 min at 4 °C on a rotating wheel. The beads were washed three times with pull-down wash buffer 1 to remove excess nucleic acids. The beads were resuspended in 250 μL of pull-down wash buffer 1 before ShTniQ (purified following Protocol 1 and thus containing co-purifying *E. coli* S15) and ShTnsC were added in 30-fold and 10-fold molar excess, respectively. ATP was added to a final concentration of 1 mM. The sample was incubated for 30 min at 37 °C, then washed three times with pull-down wash buffer 2 (20 mM HEPES-KOH pH 7.5, 250 mM KCl, 10 mM MgCl_2_, 1 mM DTT, 1 mM ATP, 0.05 % Tween20) and eluted with pull-down elution buffer (20 mM HEPES-KOH pH 7.5, 250 mM KCl, 10 mM MgCl_2_, 1 mM DTT, 1 mM ATP, 5 mM desthiobiotin). The eluted sample were analyzed by SDS-PAGE using Any kDa gradient polyacrylamide gels (Bio-Rad) stained with Coomassie Brilliant Blue and by denaturing PAGE on a 10 % polyacrylamide-7 M urea gel upon proteinase K digestion for 15 min at 37 °C prior to preparation of cryo-EM grids. Of note, recombinantly produced S15 was not added to the sample subjected to structural analysis. S15 was identified as a component of the Cas12k-transposon recruitment complex after structure determination and model building and was confirmed to have co-purified with TniQ using mass spectrometry as described above (see also **Table S2**). Prior to structural analysis by cryo-EM, complex homogeneity was assessed by negative stain electron microscopy. Negative stain EM grids were prepared as described above using a sample that was 40 times diluted as compared to the one used for cryo-EM. For preparation of cryo-EM grids, 2.5 μL of sample was applied to glow-discharged 200-mesh copper 2 nm C R1.2/1.3 cryo-EM grids (Quantifoil Micro Tools), blotted 3 s at 75 % humidity, 4 °C, plunge frozen in liquid ethane (using a Vitrobot Mark IV plunger, FEI) and stored in liquid nitrogen. Cryo-EM data collection was performed on a FEI Titan Krios G3i microscope (University of Zurich, Switzerland) operated at 300 kV equipped with a Gatan K3 direct electron detector in super-resolution counting mode. A total of 16,165 movies were recorded at a calibrated magnification of 130,000 × resulting in super-resolution pixel size of 0.325 Å. Each movie comprised 36 subframes with a total dose of 67.68 e− Å^−2^. Data acquisition was performed with EPU Automated Data Acquisition Software for Single Particle Analysis (ThermoFisher) with three shots per hole at −1.0 μm to −2.4 μm defocus (0.2 μm steps).

### Data processing and model building - Cas12k-transposon recruitment complex

Images were processed using RELION (Punjani et al., 2017) and SPHIRE (Moriya et al., 2017) software packages. All movies were motion-corrected and dose-weighted with MotionCor2(Zheng et al., 2017) (RELION). Aligned, non-dose-weighted micrographs were then used to estimate the contrast transfer function (CTF) with Gctf (Zhang, 2016). Motion corrected movies were linked to SPHIRE and corresponding image shift information converted to enable drift assessment in SPHIRE (0-60 Å overall drift selected). Single particles were picked with crYOLO (gmodel_phosnet_202005_N63_c17.h5) on JANNI denoised micrographs (gmodel_janni_20190703.h5). Particle coordinates (3.1 million) were linked back to RELION, extracted with a box size of 480 pix (binned 4-fold) and 2D classified. All particle representing classes were selected for a 3D classification using a loosely masked volume of the Cas12k-sgRNA-dsDNA complex (PDB: 7PLA, EMDB: EMD-13486 (Querques et al., 2021)) as reference to force orientational alignment on the Cas12k-sgRNA part. A 3D class showing features expanding the input density was subclassified without a mask to allow for identification of sub-populations. Sub-populations with new features were used as input (masked 3D classification) to identify all corresponding particles. Classes were inspected visually before unbinning, re-extraction and 3D refinement (RELION) in real scale. Two rounds (one round for the ‘non-productive’ complex) of Bayesian particle polishing (RELION) and CTF refinement (RELION) prior to final refinements (RELION) resulted in a reconstruction of the Cas12k-transposon recruitment complex from 75 k particles at an overall resolution of 3.3 Å (133 k particles at an overall resolution of 4.1 Å for the ‘non-productive’ complex). The local resolution was calculated based on the resulting map using the local resolution functionality (RELION) and plotted on the map using UCSF Chimera (Pettersen et al., 2004). The structure model for the Cas12k-transposon recruitment complex was built in Coot (Emsley et al., 2010). The structures of the Cas12k-sgRNA-target DNA (PDB: 7PLA (Querques et al., 2021)) and TniQ-capped TnsC filament complex (described below) were docked in the new density and used as starting model to complete the full target recognition complex. The model building revealed a well resolved extra density between tracrRNA and Cas12k. A *de novo* built template model resulting from this extra density was subjected to a DALI (Holm, 2022) search and identified the prokaryotic ribosomal S15 protein as closely related. The *E. coli* ribosomal S15 protein was confirmed to have co-purified with TniQ by mass spectrometry (see also **Table S2**) and build in the extra density with great fit. The model was refined in Coot using restraints for the nucleic acids calculated with the LibG (Brown et al., 2015) script (base pair, stacking plane and sugar pucker restraints) in ccp4 and finally refined using Phenix (Afonine et al., 2018; Liebschner et al., 2019). Real space refinement was performed with the global minimization and atomic displacement parameter (ADP) refinement options selected. Secondary structure restraints, side chain rotamer restraints, and Ramachandran restraints were used. Key refinement statistics are listed in **Table S1**. The final atomic model includes Cas12k residues 1-142, 174-636, sgRNA nucleotides 5-250, 41 nucleotides of each TS and NTS DNA, TniQ residues 9-167, S15 residues 3-87, seven TnsC molecules (each with residues 17-276), two zinc and seven magnesium cations and seven ATP molecules. The quality of the atomic model, including basic protein and DNA geometry, Ramachandran plots, clash analysis and model cross-validation, was assessed with MolProbity (Chen et al., 2010; Prisant et al., 2020) and the validation tools in Phenix. Structural superposition was performed in Coot using the secondary structure matching (Krissinel and Henrick, 2004) (SSM) function. Figure preparation for maps and models and calculation of map contour levels was performed using UCSF ChimeraX (Pettersen et al., 2021).

### cryo-EM sample preparation and data collection: TniQ-capped TnsC filament complex

For the reconstitution of TnsC filaments bound to TniQ, wild-type TnsC protein was diluted to a final concentration of 15 μM in a buffer containing 20 mM HEPES-KOH pH 7.5, 100 mM KCl, 10 mM MgCl_2_, 1 mM DTT and 1 mM ATP and mixed at a ratio of 1:25 (TnsC:DNA) with a double-stranded 69-bp duplex DNA oligonucleotide (**Table S3**) and at a 1:2 ratio (TnsC:TniQ) with TniQ in a 22.4 μl reaction volume. For preparation of cryo-EM grids, 3.0 μL of sample was applied to glow-discharged 200-mesh copper 2 nm C R1.2/1.3 cryo-EM grids (Quantifoil Micro Tools), blotted 1 s at 100 % humidity, 4 °C, plunge frozen in liquid ethane (using a Vitrobot Mark IV plunger, FEI) and stored in liquid nitrogen. Cryo-EM data collection was performed on a FEI Titan Krios G3i microscope (University of Zurich, Switzerland) operated at 300 kV equipped with a Gatan K3 direct electron detector in super-resolution counting mode. A total of 10,436 movies were recorded at a calibrated magnification of 130,000 × resulting in super-resolution pixel size of 0.325 Å. Each movie comprised 36 subframes with a total dose of 66.036 e− Å^−2^. Data acquisition was performed with EPU Automated Data Acquisition Software for Single Particle Analysis (ThermoFisher) with three shots per hole at −1.0 μm to −2.4 μm defocus (0.2 μm steps).

### Data processing and model building: TniQ-capped TnsC filament complex

Data was processed using cryoSPARC (Punjani et al., 2017). 10,436 movies were imported and motion-corrected (patch motion correction (multi)). Upon patch CTF estimation (multi), exposures were selected by estimated resolution (better than 3.5 Å) and defocus (<2 μm), yielding 10,176 movies. Particles with a particle diameter between 100 Å and 150 Å were picked on denoised micrographs using the blob picker function in cryoSPARC (Punjani et al., 2017). Particle picks were inspected and selected particles (974,175) extracted in a box size of 1200 × 1200 pixel (fourier cropped to 300 × 300 pixel) and subjected to 2D classification. Using such a large box size allowed to distinguish between classes containing shorter filaments that could be treated with a single particle cryo-EM approach from longer, continuous filaments. Classes visualizing shorter filaments (up to three rings) were selected (5 classes, 40,689) and particles were re-extracted in a box size of 600 × 600 pixel (fourier cropped to 300 × 300 pixel) and the particle stack was used to calculate two *ab initio* models. One *ab initio* models matching the appearance of 2D classes was selected as volume for non-uniform refinement. Particles were re-extracted in unbinned form and subjected to non-uniform refinement using the selected size-corrected *ab initio* volume as input. The use of cryoSPARC 3D variability analysis allowed to identify one class of two where stronger density for TniQ was visible. The corresponding particle stack (35,964 particles) was subjected to non-uniform refinement, yielding a final cryo-EM reconstruction at an overall resolution of 3.44 Å. The local resolution was calculated based on the resulting map using the local resolution functionality in cryoSPARC (Punjani et al., 2017) and plotted on the map using UCSF Chimera (Pettersen et al., 2004).

For model building, the crystal structure of TniQ (PDB:7OXD (Querques et al., 2021)) and single TnsC protomers in the cryo-EM structure of the DNA- and AMPPNP-bound TnsC filament (PDB:7PLH and EMDB: EMD-13489 (Querques et al., 2021)) were manually docked as rigid bodies in the cryo-EM density map of the TniQ-capped TnsC filament complex with UCSF Chimera (Pettersen et al., 2004), followed by real space fitting with the Fit in Map function. The DNA duplex was manually built in Coot and then subjected to refinement in Coot (Emsley et al., 2010) using base pair, stacking plane and sugar pucker restraints generated in LibG (Brown et al., 2015) in ccp4. The sequence of the dsDNA was randomly selected. The model was finally refined using Phenix (Afonine et al., 2018; Liebschner et al., 2019). Real space refinement was performed with the global minimization and atomic displacement parameter (ADP) refinement options selected. Secondary structure restraints, side chain rotamer restraints, and Ramachandran restraints were used. Key refinement statistics are listed in **Table S1**. The final atomic model includes 3 copies of TniQ, 7 copies of TnsC, each bound to a single Mg^2+^ ion and ATP. The quality of the atomic model, including basic protein and DNA geometry, Ramachandran plots, and clash analysis, was assessed and validated with MolProbity (Chen et al., 2010; Prisant et al., 2020) and validation tools as implemented in Phenix. Structural superposition was performed in Coot using the secondary structure matching (Krissinel and Henrick, 2004) (SSM) function. Figure preparation for maps and models was performed using UCSF ChimeraX (Pettersen et al., 2021).

### Transposition assays and droplet digital PCR analysis

Transposition assay were conducted as in (Saito et al., 2021), with minor modifications. All *in vivo* transposition experiments were performed in One Shot PIR1 *E. coli* cells (Thermo Fisher Scientific). The strain was first co-transformed with 20 ng each of pDonor and pTarget, and transformants were isolated by selective plating on double antibiotic LB-agar plates. Liquid cultures were then inoculated from single colonies, and the resulting strains were made chemically competent using standard methods, aliquoted and snap frozen. pHelper plasmids (20 ng) were then introduced in a new transformation reaction by heat shock, and after recovering cells in fresh LB medium at 37 °C for 1 h, cells were plated on triple antibiotic LB-agar plates containing 100 μg mL^−1^ carbenicillin, 50 μg mL^−1^ kanamycin, and 33 μg mL^−1^ chloramphenicol. After overnight growth at 37 °C for 16 h, colonies were harvested from the plates, resuspended in 15 μL Lysis buffer (TE with 0.1 % Triton X 100) and heated for 5 min at 95 °C. 60 μL of water were added to the samples before centrifugation for 10 min at 16,000 x *g*. The supernatant was then transferred and the nucleic acid concentration adjusted to 0.3 ng μL^-1^. 0.75 ng of template DNA were used for subsequent investigation by droplet digital PCR (ddPCR).

*In vitro* transposition assays were conducted using individually purified Cas12k, TnsC, TniQ, TnsB, EcS15 and ShS15 proteins, sgRNA and pTarget and pDonor plasmids (Saito et al., 2021) in 20 μL reaction volume per time point. Cas12k, TnsC, TniQ, TnsB, EcS15 and ShS15 were diluted with dilution buffer (25 mM Tris pH 8.0, 500 mM NaCl, 1 mM EDTA, 1 mM DTT, 25 % glycerol) to a concentration of 2 μM, 2.8 μM, 2.0 μM, 2.4 μM, 224 μM and 120 μM respectively. A 2X transposition buffer (50 mM HEPES-KOH pH 7.5, 40 mM KCl, 4 mM DTT, 100 μg/mL BSA, 4 mM ATP, 0.4 mM MgCl_2_) was mixed with water, pTarget (final concentration 1 ng/μl) and pDonor (final concentration 3.9 ng/μl) plasmids, Cas12k and/or sgRNA and incubated 10 min at 37 °C. TniQ, TnsC and TnsB, and/or Ec/ShS15 were added and samples were incubated for 7 min at 37 °C before starting the reaction by addition of a final concentration 15 mM magnesium acetate. The final concentration of pTarget was 1 ng/μl (56 nM), further components were added in a molar ratio of 1:5:5:60:75:5:5:5 (pTarget:pDonor:Cas12k:sgRNA:S15:TniQ:TnsC:TnsB). At respective time points, samples were heat-inactivated by incubation at 95 °C for 5 min and diluted 1/125 with dH_2_O prior ddPCR measurement.

### Pull-down experiments

For pull-down experiments using ShCas12k as bait, sgRNA was first mixed with hexahistidine-StrepII-tagged ShCas12k (Strep-Cas12k) in assembly buffer (20 mM HEPES-KOH pH 7.5, 250 mM KCl, 10 mM MgCl_2_, 1 mM DTT), and incubated 20 min at RT to allow complex formation. A dsDNA target (**Table S3**) was then added to the reaction in a Strep-Cas12k:sgRNA:dsDNA molar ratio of 1:1.2:1.5 and incubated for 20 min at RT. The final 20 μL reaction contained 5 μg (at a concentration of 3.2 μM) Strep-Cas12k, 3.8 μM sgRNA, and 4.7 μM DNA in assembly buffer. Samples were mixed with 12.5 μL Strep-Tactin beads (iba) equilibrated in pull-down wash buffer 1 (20 mM HEPES-KOH pH 7.5, 250 mM KCl, 10 mM MgCl_2_, 1 mM DTT, 0.05 % Tween20) and incubated 30 min at 4 °C on a rotating wheel. The beads were washed three times with pull-down wash buffer 2 (20 mM HEPES-KOH pH 7.5, 250 mM KCl, 10 mM MgCl_2_, 1 mM DTT, 1 mM ATP, 0.05 % Tween20) to remove excess nucleic acids. The beads were resuspended in 150 μL of pull-down wash buffer 2 and EcS15 and/or ShS15 and/or ShTniQ (purified according to Protocol 2 and thus S15-free) was added in 10-fold molar excess and/or ShTnsC was added in 12-fold molar excess.

Samples were incubated 20 min at RT, then washed three times with pull-down wash buffer 2 and eluted with pull-down elution buffer (20 mM HEPES-KOH pH 7.5, 250 mM KCl, 10 mM MgCl_2_, 1 mM DTT, 1 mM ATP, 5 mM desthiobiotin). Eluted samples were analyzed by SDS-PAGE using Any kDa gradient polyacrylamide gels (Bio-Rad) and stained with Coomassie Brilliant Blue.

## Acknowledgements

We thank Susanne Kreutzer and the ETH Zurich Genome Engineering and Measurement Lab for access to ddPCR instrumentation. We thank the Functional Genomics Center Zurich for performing the mass spectrometry analyses. This work was supported by Swiss National Science Foundation Project Grant 31003A_182567 and European Research Council (ERC) Consolidator Grant no. ERC-CoG-820152. M.S. is a member of the Biomolecular Structure and Mechanism PhD Program of the Life Science Zurich Graduate School. I.Q. was supported by FEBS and EMBO (ALTF 296-2020) long-term postdoctoral fellowships. M.J. is an International Research Scholar of the Howard Hughes Medical Institute, and Vallee Scholar of the Bert L & N Kuggie Vallee Foundation.

## Author contributions

M.S., I.Q., and M.J. conceived the study and designed experiments. M.S. prepared cryo-EM samples, collected cryo-EM data and determined the reconstruction of the Cas12k-TnsC transposon recruitment complex. M.S, I.Q. and M.J. built the model and performed structural analysis of the Cas12k-TnsC transposon recruitment complex. I.Q. prepared cryo-EM samples, collected cryo-EM data and determined the structure of the TnsC-TniQ capped filament. M.S., I.Q. and S.O. carried out biochemical and ddPCR functional experiments and negative-stain EM analysis. S.O. and C.C. assisted with sample preparation for biochemical and ddPCR assays. M.S., I.Q. and M.J. analyzed the data and wrote the manuscript.

